# PreQual: An automated pipeline for integrated preprocessing and quality assurance of diffusion weighted MRI images

**DOI:** 10.1101/2020.09.14.260240

**Authors:** Leon Y. Cai, Qi Yang, Colin B. Hansen, Vishwesh Nath, Karthik Ramadass, Graham W. Johnson, Benjamin N. Conrad, Brian D. Boyd, John P. Begnoche, Lori L. Beason-Held, Andrea T. Shafer, Susan M. Resnick, Warren D. Taylor, Gavin R. Price, Victoria L. Morgan, Baxter P. Rogers, Kurt G. Schilling, Bennett A. Landman

## Abstract

**Purpose:** Diffusion weighted MRI imaging (DWI) is often subject to low signal-to-noise ratios (SNRs) and artifacts. Recent work has produced software tools that can correct individual problems, but these tools have not been combined with each other and with quality assurance (QA). A single integrated pipeline is proposed to perform DWI preprocessing with a spectrum of tools and produce an intuitive QA document.

**Methods:** The proposed pipeline, built around the FSL, MRTrix3, and ANTs software packages, performs DWI denoising; inter-scan intensity normalization; susceptibility-, eddy current-, and motion-induced artifact correction; and slice-wise signal drop-out imputation. To perform QA on the raw and preprocessed data and each preprocessing operation, the pipeline documents qualitative visualizations, quantitative plots, gradient verifications, and tensor goodness-of-fit and fractional anisotropy analyses.

**Results:** Raw DWI data were preprocessed and quality checked with the proposed pipeline and demonstrated improved SNRs; physiologic intensity ratios; corrected susceptibility-, eddy current-, and motion-induced artifacts; imputed signal-lost slices; and improved tensor fits. The pipeline identified incorrect gradient configurations and file-type conversion errors and was shown to be effective on externally available datasets.

**Conclusion:** The proposed pipeline is a single integrated pipeline that combines established diffusion preprocessing tools from major MRI-focused software packages with intuitive QA.

## Introduction

Diffusion weighted MRI imaging (DWI) is a powerful and noninvasive way of ascertaining the microstructural makeup of the brain (1). It forms the basis for many studies including those investigating autism (2,3), aging (4,5), multiple sclerosis (6,7), schizophrenia (8,9), and the structural human connectome (10) and has been increasingly used in context of neurosurgical planning and outcomes (11–14).

However, DWI images are often subject to low signal-to-noise ratios (SNRs) and a multitude of artifacts (15). The echo planar imaging (EPI) acquisitions normally used to acquire DWI images can result in susceptibility- and eddy current-induced distortions as well as slice-wise signal drop-out (16). Imaging the brain repeatedly to acquire volumes along different gradient directions can result in long acquisition times and increased subject movement, causing significant inter-volume motion artifacts (16). Acquisitions with “prescan” turned on can result in different gain settings within a session, causing the intensities of different DWI scans to violate their expected physical relationships. Intensity distributions of the same shell should be roughly equivalent and those of larger shells should be lower than those of smaller shells (15).

Unlike other MRI modalities, DWI image file formats must store gradient configurations (b-values and b-vectors) in addition to voxel-wise intensities and coordinate space transformations. The scanner-based DICOM file format stores gradient information in the file header, but the processing-based NIFTI file format cannot. Thus, NIFTI files require separate “bval” and “bvec” text files to store this information. Converting DICOMs to NIFTIs can corrupt the data (17). For instance, some converters will reorder the volumes by shell or b-value while others do not. This can result in a mismatch between the NIFTI volumes and the gradients in the “bval” and “bvec” text files if the latter are not also reordered.

These opportunities for distortion and corruption present significant problems for DWI analysis. Subject movement has been shown to produce phase errors (18) and affect model fitting (16,19). Noise has been identified as a source of bias for DWI microstructural scalar metrics and has been shown artificially inflate some metrics in context of diseases like acute ischemia (19,20). These artifacts have also been shown to bias tractography and subsequent structural connectivity analyses of the brain (21,22). Last, they have even been shown to reduce effect sizes and induce artificial group differences in DWI analyses (20,23).

Fortunately, previous work in the field has resulted in many powerful tools across many software packages that can help correct these issues. MRTrix3’s *dwidenoise* performs Marchenko-Pastur PCA denoising on DWI images which has been shown to improve the quality of subsequent analyses (24–26). FSL’s *topup* corrects susceptibility-induced distortions (27,28). The Synb0-DisCo deep learning framework creates a synthetic b = 0 s/mm^2^ volume of infinite bandwidth from T1 images for use with *topup* when the typically necessary complementary (i.e., forward and reverse or “blip-up and blip-down”) phase encoded DWI images are not otherwise available (29). FSL’s *eddy* corrects eddy current-induced distortions and motion artifacts, imputes slices with signal drop-out, and rotates gradients to better align them with DWI signals (30,31). Last, MRTrix3’s *dwigradcheck* determines optimal DWI gradient configurations, defined as the gradient axis order and sign permutation that yields the highest average whole-brain streamline length (32).

The existence of these tools, however, does not ensure that all barriers to robust and artifact-free diffusion analyses have been removed. For instance, because these tools are primarily used separately, manipulation of data between software packages with different syntax standards, image spaces, and formatting expectations presents an additional opportunity for data corruption. Additionally, there will inevitably be instances when algorithms underperform, and these tools fail. Thus, the use of the preprocessing tools themselves presents an opportunity for data corruption. This suggests that optimal use of these tools would be in an integrated pipeline for reproducible preprocessing that not only applies the tools to the data but also provides a way to perform quality assurance (QA) on each tool in addition to the data itself. Considering the high dimensionality of DWI data even in a single acquisition, the pipeline’s QA process would necessarily need to be intuitive but still comprehensive. Last, due to the extreme diversity of diffusion representations, models, and analyses, such a preprocessing and QA pipeline would need to be generalizable and prepare DWI data for any arbitrary secondary analysis or modeling.

Some solutions to this problem currently exist. One pipeline, built by Lauzon et al. in 2013, analyzes the quality of DWI data and details the results in an intuitive document for user review. However, it was designed prior to the release of many of these preprocessing tools and thus does not perform extensive distortion correction (33). Another pipeline, *DTIPrep*, performs distortion correction with QA, but does not support susceptibility-induced distortion correction and requires users learn to use its custom QA software in order to understand its results (34). FSL's *eddyqc* performs QA on *eddy* (35) and details the results in an easily understandable output document but does not provide integration with or QA of other tools. More recently, the DESIGNER pipeline integrates many of these tools in a simple user interface but does not provide any QA (36), and the TORTOISE pipeline implements many of its own preprocessing steps but provides limited QA (37–39). Last, the QSIPrep pipeline offers extensive preprocessing and modeling options but does not offer a robust QA of the raw images, does not take advantage of the most modern deep learning distortion correction techniques (29), and exhibits reduced generalizability by performing secondary modeling and analyses within its own framework (40).

Thus, we propose PreQual, a pipeline designed to fill this gap. By leveraging and combining powerful software tools from a variety of packages into a single pipeline, we produce preprocessed DWI outputs ready for arbitrary secondary analysis. Additionally, by calculating summary statistics, visualizing trends and image volumes, and interrogating a robust diffusion tensor imaging (DTI) model of the data, we perform dedicated qualitative and quantitative QA on each preprocessing step and on the raw input and preprocessed output data as a whole. Last, to comprehensively and intuitively report the QA results to users, the pipeline saves all intermediates and statistics for user investigation and culminates in a QA report in the common portable document format (PDF) with dedicated pages and fields for each preprocessing and QA operation.

## Methods

We built PreQual around the MRTrix3 (41), FSL (42), and ANTs (43) software packages, MRI-focused tools popular in the field, in order to perform integrated preprocessing, artifact correction, and intuitive QA (Figure 1).

**Figure 1.**
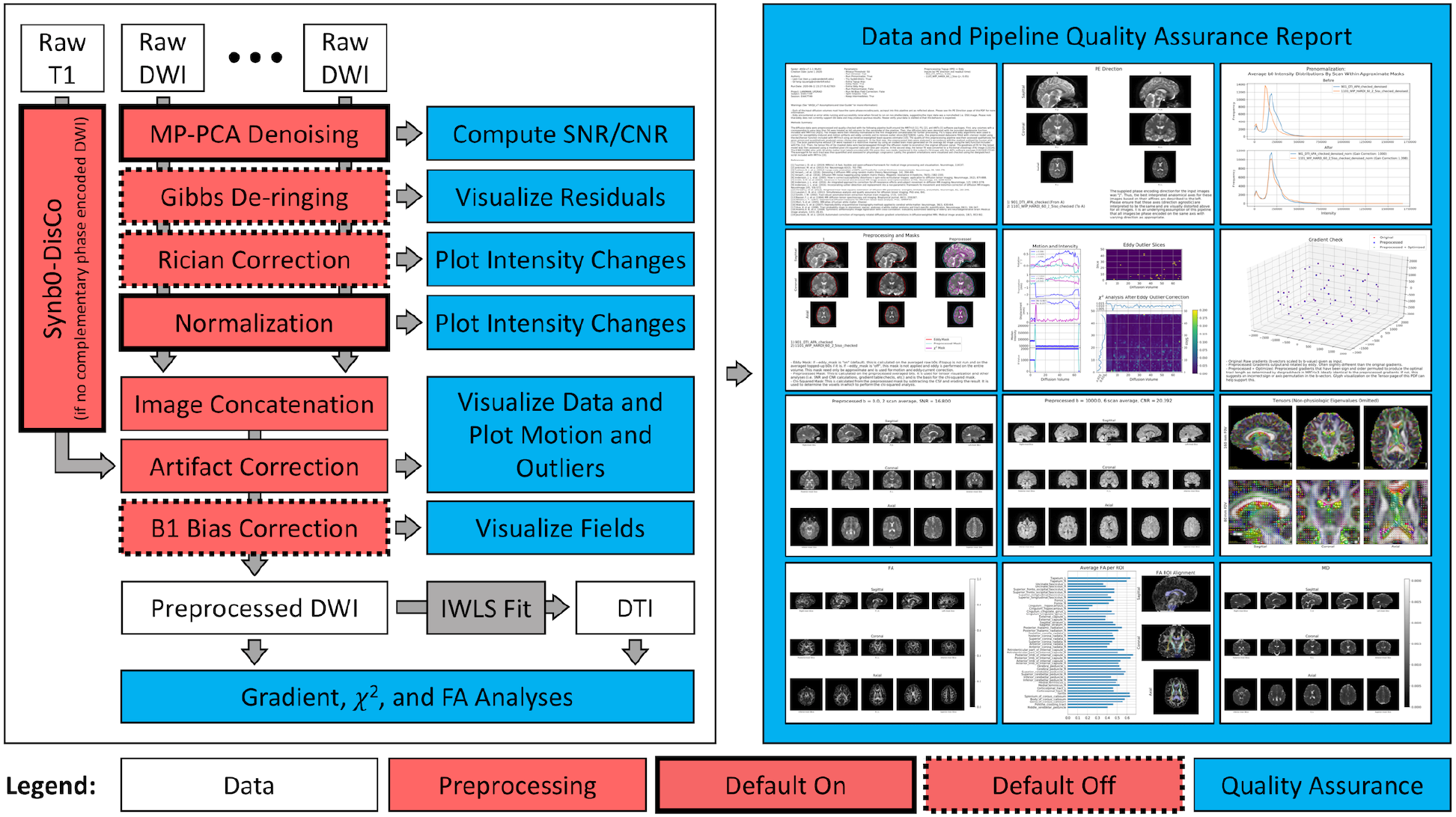
Overview of the PreQual pipeline and QA document. By default, the pipeline takes as input raw DWI data, denoises them, normalizes their intensities, concatenates them, and corrects for artifacts to produce preprocessed DWI data ready for secondary analysis. If no complementary phase encoded images are supplied, PreQual uses Synb0-DisCo and a T1 image to perform susceptibility-induced distortion correction. PreQual can also perform Gibbs de-ringing, Rician correction, and B1 bias field correction if the user wishes. QA checks on each preprocessing step are performed. The pipeline fits the preprocessed DWI data to a tensor model with an iterative weighted least squares (IWLS) estimator to produce a DTI image. Together, the DWI and DTI data are interrogated in order to QA the preprocessed DWI output of the pipeline. All QA processes are presented in the output report. All preprocessing steps except for image concatenation and artifact correction are optional and can be turned on or off per user preference.

### Raw DWI Inputs

We take a series of raw DWI images as input into the pipeline along with a phase encoding axis and a configuration file. We expect the input images as NIFTI files, properly converted from DICOM files (Supporting Information Table S1). Specifically, we expect that all gradients have been reoriented into subject space and that reordering of image volumes did not occur. We find that the most reliable way of achieving this is with recent versions of the popular *dcm2niix* tool (17). Regarding input configurations, we envision the input images to be from the same acquisition session, phase encoded along the same axis, defined as the image dimension along which phase encoding occurred. In the configuration file, we require the user to input the directions along the axis in which the images were encoded and their corresponding readout times. The phase encoding schemes of each image (i.e., axis, direction, and readout time) are required for susceptibility-induced distortion correction with FSL’s *topup* (27,28).

We note that the supplied phase encoding axis of the inputs may not always correctly correspond to the expected anatomical axis. For instance, DWI images phase encoded along the anterior to posterior axis and oriented radiologically (right-anterior-superior) or neurologically (left-anterior-superior) would be encoded along the second dimension. Yet, if the image is oriented differently and anterior to posterior phase encoding is expected, the user may need to indicate a different dimension for the phase encoding axis. Thus, for QA we check that the supplied axis and direction for each image translates to the expected anatomical axis and direction using each image’s coordinate transformation. We visualize triplanar slices of the raw data alongside these interpretations for users to verify the presence of susceptibility-induced distortions along the anatomical axes (Supporting Information Figure S1).

### Denoising

We individually denoise each input image using the Marchenko-Pastur PCA technique as implemented in MRTrix3 (24–26). For QA, we calculate the median intra-brain voxel-wise SNR for the non-diffusion weighted b = 0 s/mm^2^ volumes and display it in the QA document (Supporting Information Figure S7). We define SNR in alignment with FSL’s *eddy* (Eq. 1) as the mean voxel-wise intensity divided by its standard deviation.

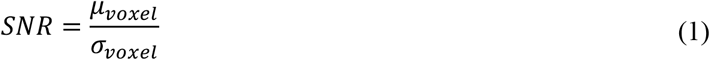

When only one b = 0 s/mm^2^ volume is available, we report the SNR as “Not a Number”. This step is on by default but can disabled by users. In this instance, we still display the SNR as a method of QA.

### Normalization

We correct for inter-scan gain differences between separate input image files that can arise when a scanner’s “prescan” setting is enabled. To achieve this, for the *k*th input image file, we take the average b = 0 s/mm^2^ volume, *V*_*k*_, and mask the result with FSL’s brain extraction tool (*bet*) (44). We then determine a multiplicative scale factor, *s*_*k*_, such that the histogram intersection, *I*, between the intra-mask intensity histogram, *h*, of *V*_*k*_ scaled by *S*_*k*_ and that of the first image is maximized. When *k* = 1, *s*_1_ = 1. When *k* > 1, we calculate *s*_*k*_ by performing optimization on the negated problem (Eq. 2) with the downhill simplex algorithm with 10 initial guesses evenly spaced between 0.5 and 1.5 (45).

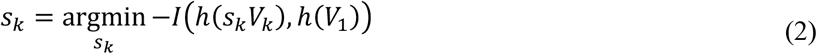

During each iteration of the algorithm, we calculate the histograms *h*(*s*_*k*_*V*_*k*_) and *h*(*V*_1_) with 1000 bins from *a* to *b* where *a* = min(*V*_1_, *s*_*k*_*V*_*k*_), the minimum intra-mask intensity between the two average b = 0 s/mm^2^ volumes, and *b* is the corresponding maximum. To complete normalization, we intensity-scale all volumes, diffusion-weighted or otherwise, of the *k*th image by *S*_*k*_ for all *k*. For QA, we report the calculated scale factors and plot of the average b = 0 s/mm^2^ intensity histograms of each image before and after normalization in the report (Supporting Information Figure S2).

We note that though prescan-induced gain changes can cause different scans to violate expected physical relationships, not all studies necessarily suffer from this. We only consider inter-scan normalization and thus do not interfere with intra-scan intensity relationships. As a result, for many multi-shell acquisitions, the intra-shell relationships are not modified because each shell is often contained in its own image file. Similarly, the inter-shell relationships in multi-shell acquisitions stored within one file are not modified. Some scanners provide explicit control for the prescan setting, so it is known that the scans already possess correct intensity relationships. In these cases, normalization could do more harm than good. Thus, we allow users to disable this step if needed, but still report the intensity histograms and the scale factors that would have been necessary for normalization for QA.

### Artifact Correction

Following normalization, we concatenate the images and perform artifact correction. First, we correct susceptibility-induced distortions using FSL’s *topup* (27,28). We generate the *topup* acquisition parameters file using the input phase encoding schemes provided and run *topup* on all the b = 0 s/mm^2^ volumes from all input scans. Typically, *topup* requires complementary phase encoded images. In the event that the user does not supply these, we use Synb0-DisCo to create a susceptibility-corrected synthetic b = 0 s/mm^2^ volume of infinite bandwidth for *topup* instead (29). When we run Synb0-DisCo, we require that the user input a T1 image of the same subject into the pipeline and visualize triplanar views of the T1 and synthetic b = 0 s/mm^2^ volume in the output QA document (Supporting Information Figure S3).

Following *topup*, we use FSL’s *eddy* with a brain mask estimated by *bet* on the averaged *topup* output to correct for eddy current-induced distortions, inter-volume motion, and slice-wise signal drop-out (30,31). During this process, *eddy* also performs rotation of the gradients to best suit the DWI signals and is configured to estimate the voxel-wise SNR (Eq. 1) across b = 0 s/mm^2^ volumes and contrast-to-noise ratio (CNR) for each diffusion weighted shell. *Eddy* defines CNR as *σ*_*GP*_/*σ*_*residual*_ or the voxel-wise standard deviation of the Gaussian process predicted by *eddy* divided by that of the residuals. A final brain mask is calculated with *bet* (44). This results in the pipeline’s final preprocessed DWI outputs.

For QA of the artifact correction steps, we take a combined quantitative and qualitative approach. First, we use the final mask to calculate the median intra-mask SNR for b = 0 s/mm^2^ volumes and CNR for each diffusion weighted shell. We display these ratios alongside five central triplanar slices of each corresponding shell-wise averaged preprocessed volume (Supporting Information Figure S7). This provides a quantitative metric for SNR for the denoising step as well as qualitative verification of artifact correction. Additionally, we plot the *eddy*-calculated inter-volume rotation, translation, and displacement and print their average values alongside a plot indicating the slices *eddy* imputed due to signal drop-out (Supporting Information Figure S5).

### Tensor Fitting

We fit the preprocessed DWI volumes to a tensor model using the iterative weighted least squares (IWLS) estimator in MRTrix3 (46) and compute fractional anisotropy (FA) and mean diffusivity (MD) scalar maps (47). As DTI is not the most appropriate model for all DWI studies, we use the tensor data and scalar maps primarily to QA the preprocessed DWI data, but we make them available for secondary analysis if the user wishes. For QA, we visualize the tensor glyphs (Supporting Information Figure S8) as rendered in MRTrix3 (48) and five central triplanar slices of the FA and MD maps in the output document (Supporting Information Figure S9).

### Gradient Analysis

To see whether the raw DWI gradient directions are properly oriented, we determine their optimality. We define optimality as it is defined in MRTrix3: the permutation of gradient sign and axis order that produces the highest average whole-brain streamline length (32). Since this process depends on tractography and tractography is affected by distortions (21,22), we use the preprocessed DWI gradient directions output by *eddy* as surrogates for the raw gradients. This substitution will not impact the analysis because although *eddy* rotates gradients, it only performs minor adjustments and does not change the gradient sign or axis order. We plot the optimal preprocessed gradient permutation, the preprocessed gradient directions as output by *eddy*, and the original raw input gradient directions (Supporting Information Figure S6) in the output QA document. We also visualize triplanar slices of the tensor glyphs (Supporting Information Figure S8). We expect properly oriented gradients to be identifiable with this visualization, as the preprocessed and optimal gradients would overlap and produce physiologically oriented tensors. Additionally, plotting of the raw gradients provides an extra check that the preprocessed gradients contain only minor deviations and are acceptable surrogates for this analysis.

Many problems can result in improperly oriented gradients. As a result, we save the optimal gradients for user analysis, but we do not overwrite the gradients output by *eddy* in the outputs of the pipeline. The rationale for this is that we have empirically found that improperly oriented gradients are rarely the result of a pure sign or axis flip but are often due to a rotational bias and thus do not wish to confuse users into believing gradient errors have been completely resolved. If problems are identified, the input data need to be corrected and reprocessed.

### Fractional Anisotropy Analysis

To quality check the preprocessed DWI data, we interrogate the calculated DTI model and compute the average FA value for all 48 white matter regions of interest (ROIs) defined by the Johns Hopkins ICBM DTI 81 white matter atlas (49–51). We deformably register the atlas to the subject’s FA image space using the ANTs software package (52) to isolate each region and its average FA value. For QA, we plot these values along with an overlay of the atlas on the subject’s FA map. The former allows users to verify regional FA congruence with expected physiologic values, and the latter allows users to check the registration process (Supporting Information Figure S10). As a heuristic, we expect the regional FA values to range from roughly 0.3 to 0.7 (53).

### Chi-Squared Analysis

To investigate the adequacy of the preprocessed DWI data for secondary modeling, we measure the goodness-of-fit of the tensor model with a modified pixel-wise chi-squared analysis as described by Lauzon et al. on the preprocessed DTI data (33). We constrain this analysis to the brain parenchyma in order to avoid analyzing tensors fit to background or CSF that would produce invalid chi-squared values. To mask the parenchyma, we erode the final mask calculated on the preprocessed DWI data with a radius of one voxel and remove the CSF. To identify the CSF, we calculate a CSF mask for each shell-wise averaged volume of the preprocessed DWI data with FSL’s *fast* tool (54). We then generate an overall CSF probability mask by performing a voxel-wise average across shells and setting a threshold at 15%. For each slice in each volume, we divide the intra-parenchyma sum squared error of tensor fit by the sum squared intensity of the slice to calculate the goodness-of-fit (33). For QA, we visualize the chi-squared values in the QA document (Supporting Information Figure S5).

### Mask Quality Assurance

We quality check the mask used for *eddy*, the final preprocessed mask used for median SNR and CNR calculation, and the parenchyma mask used for the chi-squared analysis. We plot their contours on triplanar views of the first b = 0 s/mm^2^ volume of each raw input image and on that of the preprocessed DWI output (Supporting Information Figure S4).

### Additional Pipeline Options and Considerations

For preprocessing, we also include Gibbs de-ringing with the local subvoxel-shifts method after denoising (55), Rician correction with the method of moments that does not require phase data before intensity normalization (36,56), and N4 B1 bias field correction after artifact correction (57). Gibbs rings are artifacts located adjacent and parallel to high contrast interfaces due to the truncation of Fourier domain signals when reconstructing MRI images (55). Thus, to QA Gibbs de-ringing, we visualize triplanar slices of the averaged differences across all b = 0 s/mm^2^ volumes before and after de-ringing looking for larger differences at high contrast interfaces (Supporting Information Figure S11). The construction of magnitude images from complex raw MRI data can result in noise taking on a Rician profile with positive bias, as opposed to a Gaussian one, especially with lower SNR as found in higher shells (36,58). Thus, to QA Rician correction, we plot the shell-wise intensity distributions within the brain before and after correction for each input image looking for decreased signal intensities with correction (Supporting Information Figure S11). B1 bias fields are low frequency spatially distributed intensity inhomogeneities that affect some MRI acquisitions. To QA N4 correction of these fields, we visualize triplanar slices of the estimated field and of the first b = 0 s/mm^2^ volume before and after correction (Supporting Information Figure S11). Additionally, we make running *eddy* without a brain mask an option because the brain extraction process can be unstable at times. Empirically we have found this produces minor differences in preprocessed outputs but allows preprocessing to generalize to cases with poor masking.

For QA, tensor visualization can be difficult due to crowding of glyphs for high-resolution or up-sampled data. As such, we include the ability to visualize principal eigenvectors (Supporting Information Figure S11) instead of full tensors. While losing some information about the tensor fit, this allows users to still visualize gradient direction information. Last, in addition to displaying all summary statistics (SNR and CNR; average intervolume rotation, translation, and displacement; and regional FA) in the QA report (Supporting Information Figures S5, S7, and S10), we tabulate them in a comma separated value (CSV) file so that users can extract the information as needed (Supporting Information Figure S12).

### Study Overview: Characterization of Data

We evaluate pipeline efficacy by running it on one of two DWI datasets acquired on a Philips 3T scanner at Vanderbilt University. The first dataset, Dataset A, was acquired on one subject scanned repeatedly on the same scanner over the course of three sessions, one session each consecutive day (59). Each session consisted of 6 sets of scans. Each set of scans consisted of one 3-direction b = 1000 s/mm^2^ image phase encoded in the anterior to posterior direction (APA), one 96-direction b = 1000 s/mm^2^ image phase encoded in the posterior to anterior direction (APP), one 96-direction b = 1500 s/mm^2^ APP image, one 96-direction b = 2000 s/mm^2^ APP image, one 96-direction b = 2500 s/mm^2^ APP image, and one 96-direction b = 3000 s/mm^2^ APP image. The second dataset, Dataset B, was taken from another subject scanned once. It consists of a 6-direction b = 1000 s/mm^2^ APA image and one 60-direction b = 2000 s/mm^2^ APP image. All images were acquired with corresponding b = 0 s/mm^2^ volumes and were deidentified and acquired only after informed consent under supervision of the project Institutional Review Board.

### Study Overview: Characterization of Pipeline

We study the denoising step by calculating changes in SNR of DWI data preprocessed with and without denoising. We examine the normalization step by simulating inter-scan gain differences and observing how the pipeline corrects them. We investigate the artifact correction step by visualizing representative volumes before and after correction. We explore the gradient analysis by negating input gradients in the second dimension and visualizing how the pipeline identifies this simulated error. We demonstrate one use case of the FA analysis by shuffling the order of the input image volumes without shuffling their corresponding gradients to simulate a failed file-type conversion and observing how the pipeline identifies this. We leverage the chi-squared analysis to understand how the pipeline prepares DWI data for secondary modeling by performing the same tensor fit on the raw input data as is performed on the outputs and measuring the differences in goodness-of-fit.

We study the generalizability of PreQual by running it on one imaging session from each of three external datasets that contain isolated QA issues and investigate how the pipeline identifies them. The first is a session from the Human Connectome Project (HCP) Lifespan cohort (60). We have previously identified obvious susceptibility-induced distortions in this session (29). The second is a session from the Autism Brain Imaging Data Exchange (ABIDE) II (2). We have previously identified motion-induced artifacts in this session. The third is a session from the Baltimore Longitudinal Study of Aging (BLSA) (61,62). We have previously identified DICOM to NIFTI conversion errors in this particular session that have resulted in improperly oriented gradients that have since been corrected (63). Last, we quantify the runtime of PreQual on three different configurations of Dataset B.

For this study, we perform paired statistical comparisons without the assumption of normality. Thus, we use the non-parametric Wilcoxon signed-rank test for paired distributions to determine statistical significance and report all p-values without multiple comparisons correction (64).

## Results

### Denoising

To study the impact of the denoising step, we ran PreQual on each session of Dataset A and combined the results for analysis. We calculated the median intra-mask pixel-wise SNR for each shell and display them in Figure 2. Qualitative visualization showed improved SNR for each diffusion weighted shell which were each shown to be statistically significant at p < 0.001 (Wilcoxon signed rank test).

**Figure 2.**
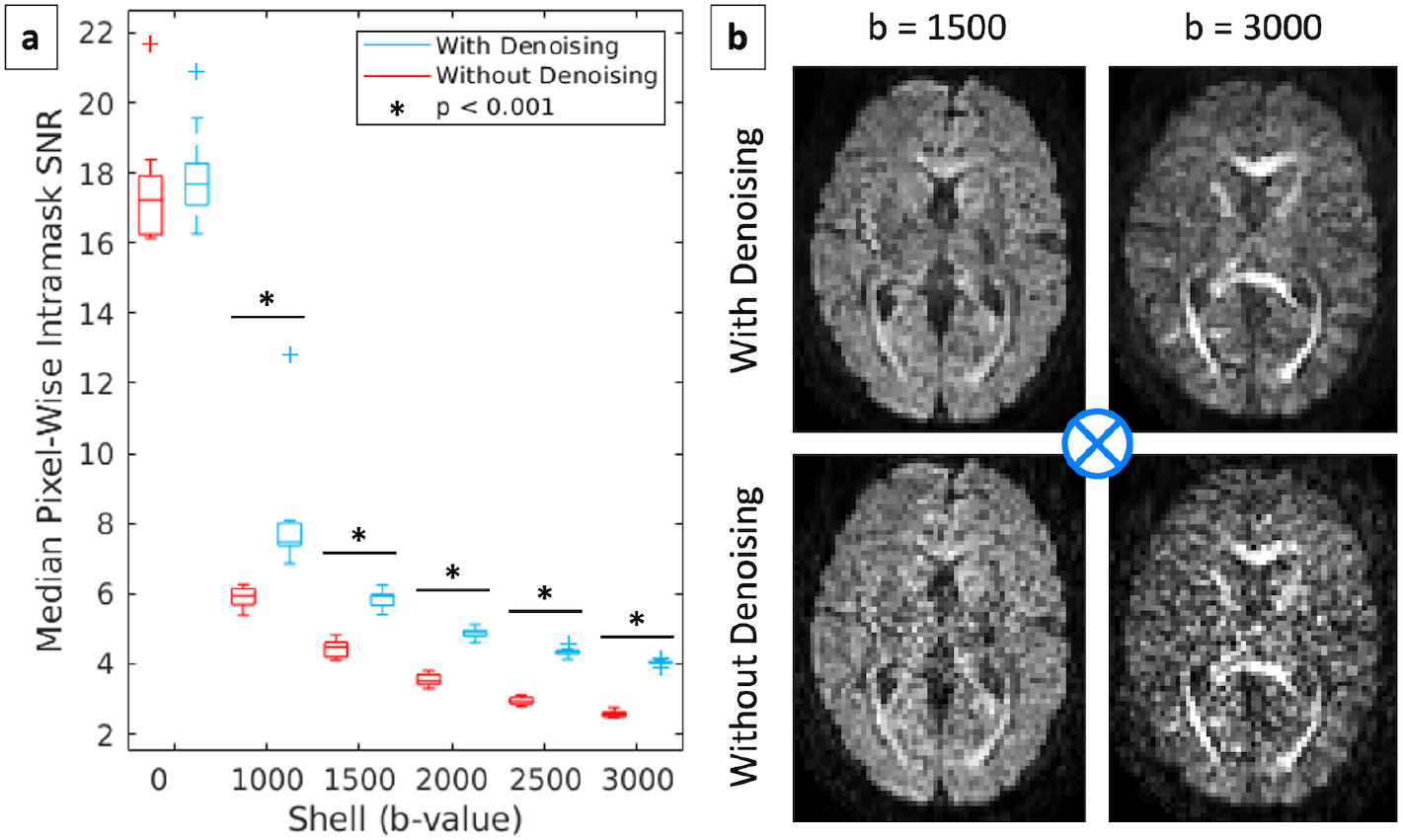
PreQual improves the signal-to-noise ratio (SNR) of diffusion images. (a) The shell-wise SNR for all diffusion weighted shells increased in the preprocessed DWI outputs of PreQual following Marchenko-Pastur PCA denoising of the input data as implemented in MRTrix3 compared to preprocessing without denoising (Wilcoxon signed rank test, p < 0.001). (b) Visualization of representative volumes at shells b = 1500 s/mm2 and b = 3000 s/mm2 with gradient direction x = 0.002, y = 0.005, and z = 0.999 (best modeled as a unit vector going into the page as shown with the blue fletching) illustrates qualitatively improved SNR.

### Normalization

To study the efficacy of the normalization step, we ran PreQual on Dataset B while simulating inter-scan gain changes. When different images have the same gain settings, we expect intensity distributions of larger shells to be lower than those of smaller shells and those of the same shell to be roughly equivalent (15). We use each volume’s median intra-mask intensity as a representative measure for its intensity distribution, and we intentionally violate both of these expectations by multiplying the b = 2000 s/mm^2^ image of Dataset B by four. This caused the median intensities of the b = 2000 s/mm^2^ volumes to be higher than those of the b = 1000 s/mm^2^ volumes (Figure 3b). It also caused the median intensities of the b = 0 s/mm^2^ volumes to be different from each other (Figure 3a). We show that the expected relationships return after normalization with PreQual (Figures 3c and 3d).

**Figure 3.**
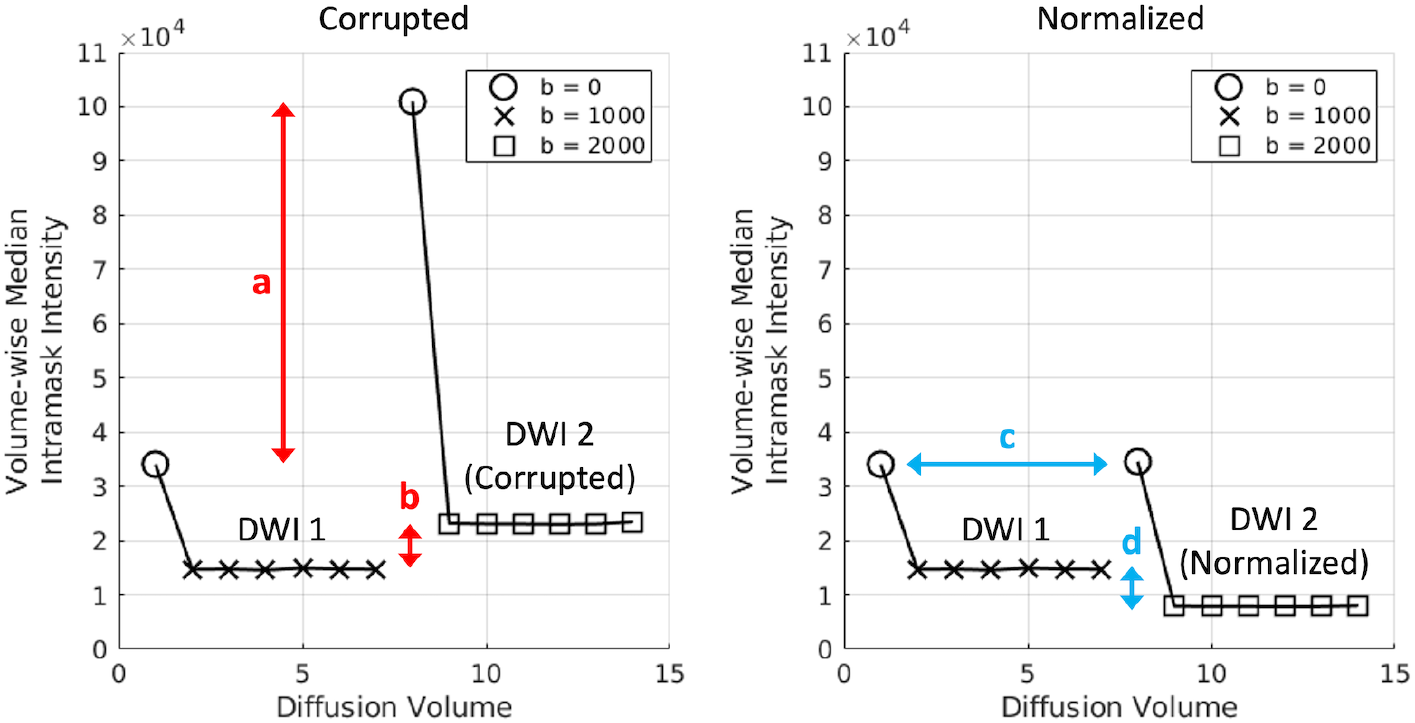
PreQual corrects physically impractical shell-wise intensity values in simulation. Two DWI images of the same session were used as input for this simulation. DWI 1 was a 6-direction b = 1000 s/mm2 image with an associated b = 0 s/mm2 volume, and DWI 2 was a 60-direction b = 2000 s/mm2 image with an associated b = 0 s/mm2 volume (only the first 6 volumes are shown for simplicity). DWI 2 was corrupted via global intensity-scaling by a factor of 4, emulating gain changes between scans of the same session. This resulted in physically impractical inter-shell intensity relationships. (a) Scans of the same shell (or b-value) are expected to have roughly the same intensity distributions, a phenomenon which is violated here in simulation. The b = 0 s/mm2 volumes are expected to have roughly the same median intensity. (b) Shell size and intensity are inversely related which is violated here in simulation. The b = 2000 s/mm2 volumes are expected to have lower median intensity than the b = 1000 s/mm2 volumes. (c and d) The expected phenomena are observed after normalization with PreQual.

### Artifact Correction

To study the effectiveness of susceptibility-, eddy current-, and motion-induced distortion correction and slice-wise signal drop-out imputation by PreQual, we ran the pipeline on both Datasets A and B and visualized representative inputs and outputs of each operation (Figures 4a, 4c, 4d, and 4e). We also ran PreQual on Dataset B with the b = 1000 s/mm^2^ APA image omitted, thus destroying the necessary complementary phase encoded pair for *topup* and triggering Synb0-DisCo to run. We visualized the inputs and outputs of this operation as well (Figure 4b). Qualitative visualization demonstrated cases where artifact correction was needed as well as the improvement of the data with PreQual.

**Figure 4.**
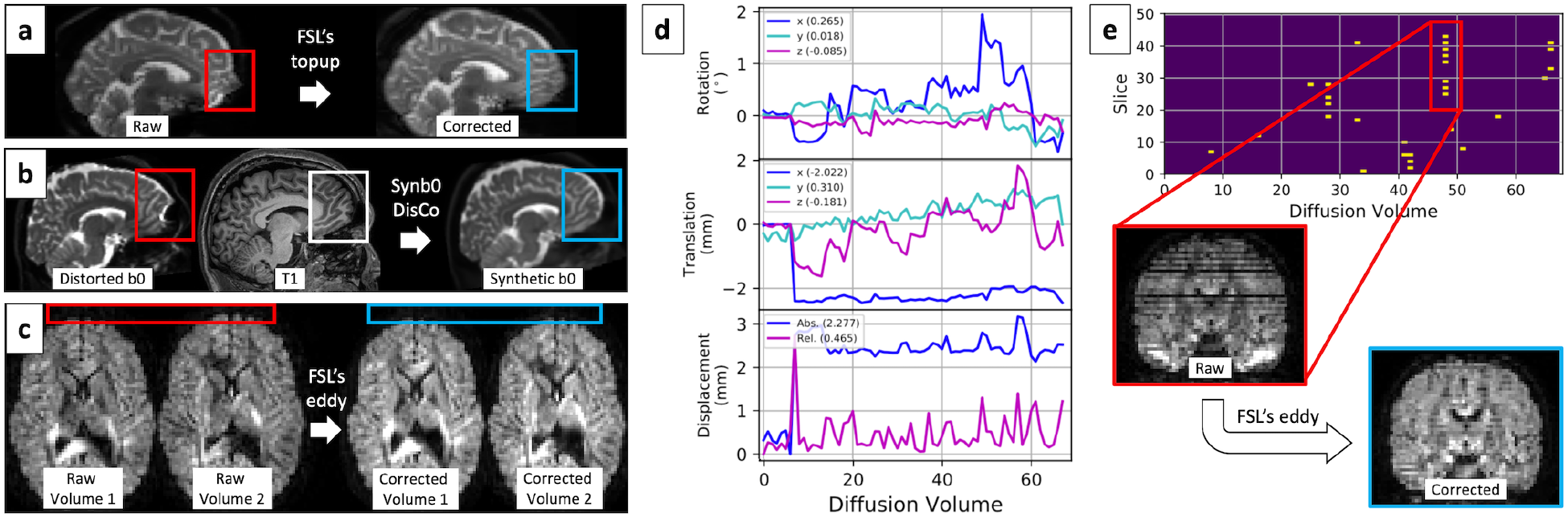
PreQual performs artifact correction. (a) Susceptibility-induced distortions are corrected using FSL’s *topup* in conjunction with input complementary phase encoded images. Examples of this distortion before and after correction with *topup* are shown in the red and blue bounding boxes, respectively. (b) If no complementary phase encoded images are available, PreQual uses Synb0-DisCo to generate a synthetic distortion-free b = 0 s/mm2 volume from a distorted one and a T1 image. The synthetic volume is used with *topup* to correct the image as a whole. The bounding boxes highlight an example of how Synb0-DisCo generates distortion-free regions (blue) by using distorted b = 0 s/mm2 information (red) and distortion-free T1 information (white). (c) Eddy current-induced inter-volume geometric distortions as highlighted between raw volumes 1 and 2 (red) are corrected with FSL’s *eddy* as shown between corrected volumes 1 and 2 (blue). (d) FSL’s *eddy* estimates and corrects for inter-volume motion. An example output in the PreQual QA report detailing this movement is shown. The overall average movement across volumes is shown in the legend parentheses. For displacement, legend entry “Abs.” is short for absolute, and “Rel.” is short for relative. (e) *Eddy* identifies slice-wise signal drop-out and imputes lost slices accordingly. An example output plot in the PreQual report details the affected slices. The corresponding raw volume with drop-out slices is shown highlighted in red and the imputed output of *eddy* is shown highlighted in blue. Note that the imputation improves data quality, but that the overall process is still quite motion-sensitive, explaining some of the remaining artifacts.

### Gradient Analysis

We ran PreQual on Dataset B in two configurations. In the first configuration, the data were not corrupted. In the second, all gradients were negated in the second dimension. We visualize the resultant tensor glyphs in the splenium of the corpus callosum for both cases and show them to be oriented in physiologically probable and improbable orientations, respectively. We also visualize the optimal gradient tables as determined by PreQual for both cases and show them to be overlapping and non-overlapping with those output in the preprocessed DWI data, respectively. These two visualizations together allow us to correctly identify the case with improperly oriented gradients (Figure 5).

**Figure 5.**
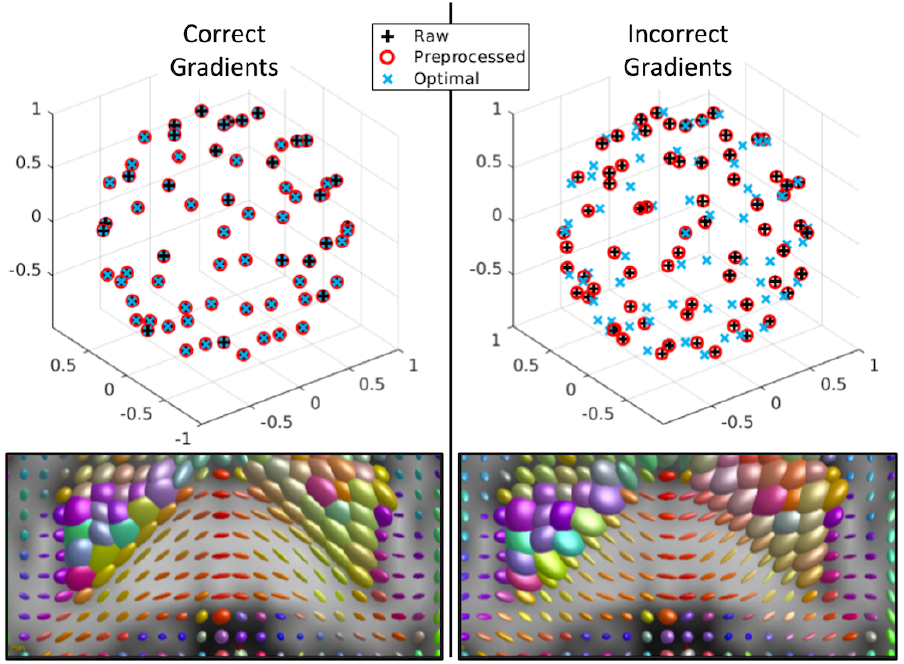
PreQual identifies improperly oriented gradient tables in simulation. The left side of the figure displays expected phenomenon. The optimal gradient configurations are identical to those in the preprocessed DWI output of PreQual. The unit gradient vectors are overlapping, and the tensors calculated from the preprocessed gradients are oriented in a physiologically probable manner, as shown in an axial slice through the splenium of the corpus callosum. The right side of the figure displays the identification of improperly oriented gradients. The preprocessed gradients do not overlap with the optimal ones and the tensor glyphs are oriented along the corpus callosum in a physiologically improbable manner. The raw input gradients are shown here to match what is reported in the QA report such that users can visualize rotational optimization made to input gradients during preprocessing.

### Fractional Anisotropy Analysis

To demonstrate one use case for the FA analysis, we simulated a failed file-type conversion that led to the mismatch between image volumes and gradient tables. We took the b = 0 s/mm^2^ volume acquired at the beginning of the b = 2000 s/mm^2^ APP scan of Dataset B out of its original position and placed in the middle of the sequence without reordering the corresponding gradients. We demonstrate that the average ROI-based FA exhibit an entirely different cross-region profile with this volume-to-gradient mismatch than without (Figure 6a). This is supported by qualitative visualization of the FA maps (Figure 6b) and together result in the identification of faulty file-type conversions.

**Figure 6.**
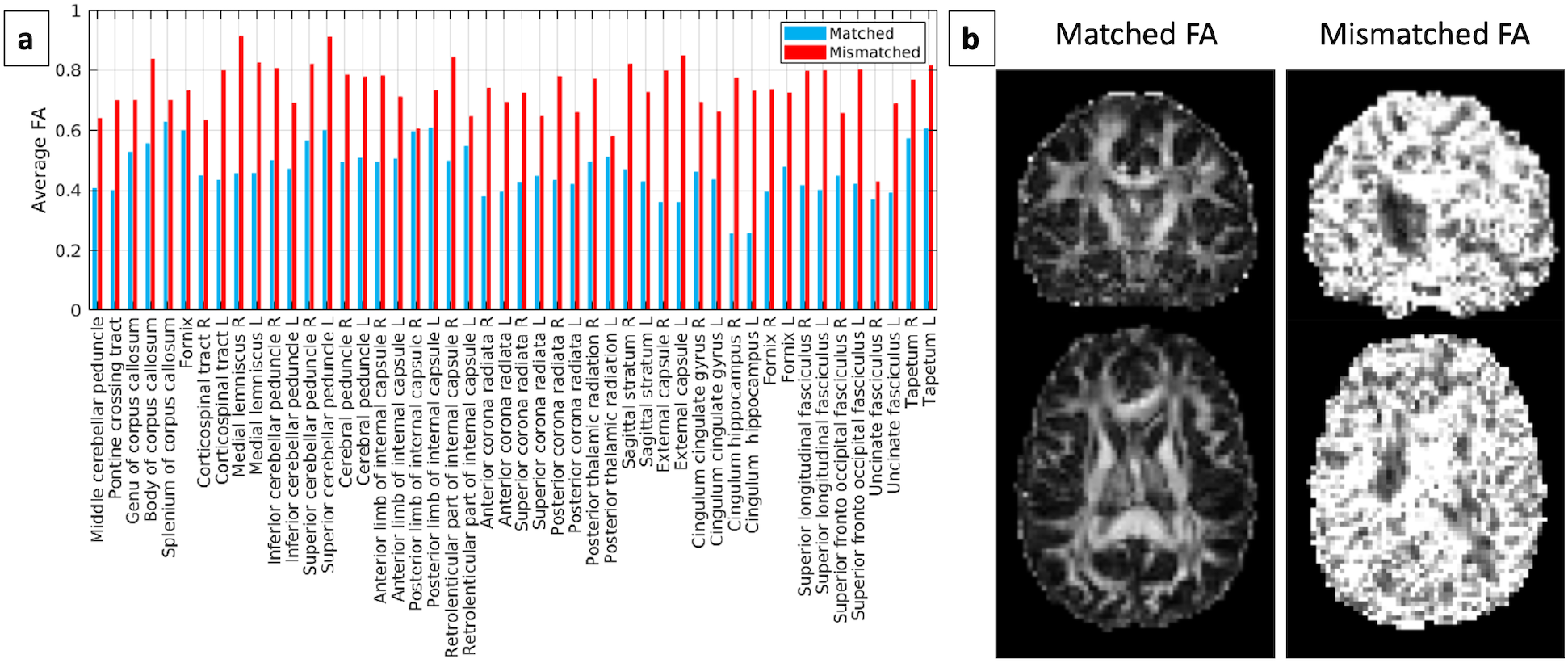
PreQual identifies simulated faulty file-type conversions that lead to volume-to-gradient mismatch. Volume-to-gradient mismatch was achieved by shuffling b = 0 s/mm2 volumes out of their original position in the DWI sequence without corresponding shuffling of gradient information. (a) The average FAs by white matter region are higher across all regions and exhibit an identifiably different cross-region profile with volume-to-gradient mismatch. (b) Qualitative examination of FA maps with and without mismatch provides additional verification.

### Chi-Squared Analysis

We show a chi-squared goodness-of-fit analysis in Figure 7. We performed the analysis on Dataset B and measured the chi-squared values calculated on the raw images and on those preprocessed with PreQual. Since chi-squared is a measurement of error, the lower values in the data preprocessed with PreQual suggest improved tensor fitting of the data (p < 0.001, Wilcoxon signed rank test).

**Figure 7.**
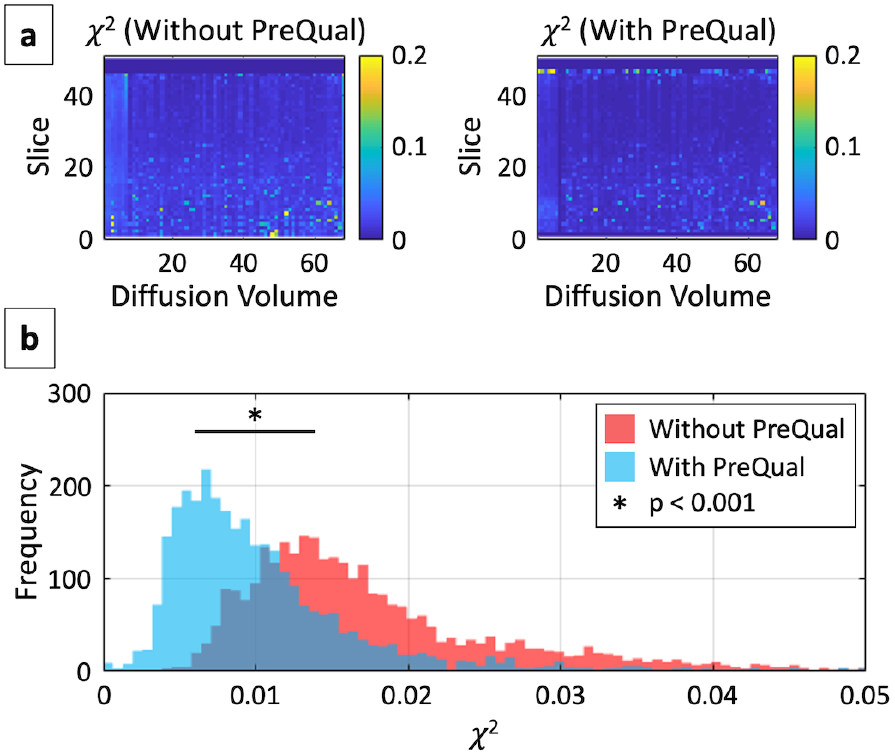
PreQual improves tensor fitting of diffusion weighted images. (a) The modified pixel chi-squared for a multi-shell acquisition with and without PreQual applied are presented per slice per volume. (b) The distributions of the chi-squared values in (a) are presented and demonstrate improved goodness-of-fit with statistical significance (Wilcoxon signed rank test, p < 0.001).

### External Verification

We ran the pipeline on one imaging session from each of three externally available datasets, as shown in Figure 8, to demonstrate the generalizability of PreQual. We show visually apparent susceptibility-induced distortions corrected in the HCP Lifespan image (60), slice-wise signal drop-out imputed in the ABIDE II image (2), and improperly oriented gradients identified in the uncorrected BLSA image (61–63). The specific sessions used are detailed in Supporting Information Table S2, and the full QA documents generated by PreQual corresponding to these datasets are available in Supporting Information Figures S1 through S10.

**Figure 8.**
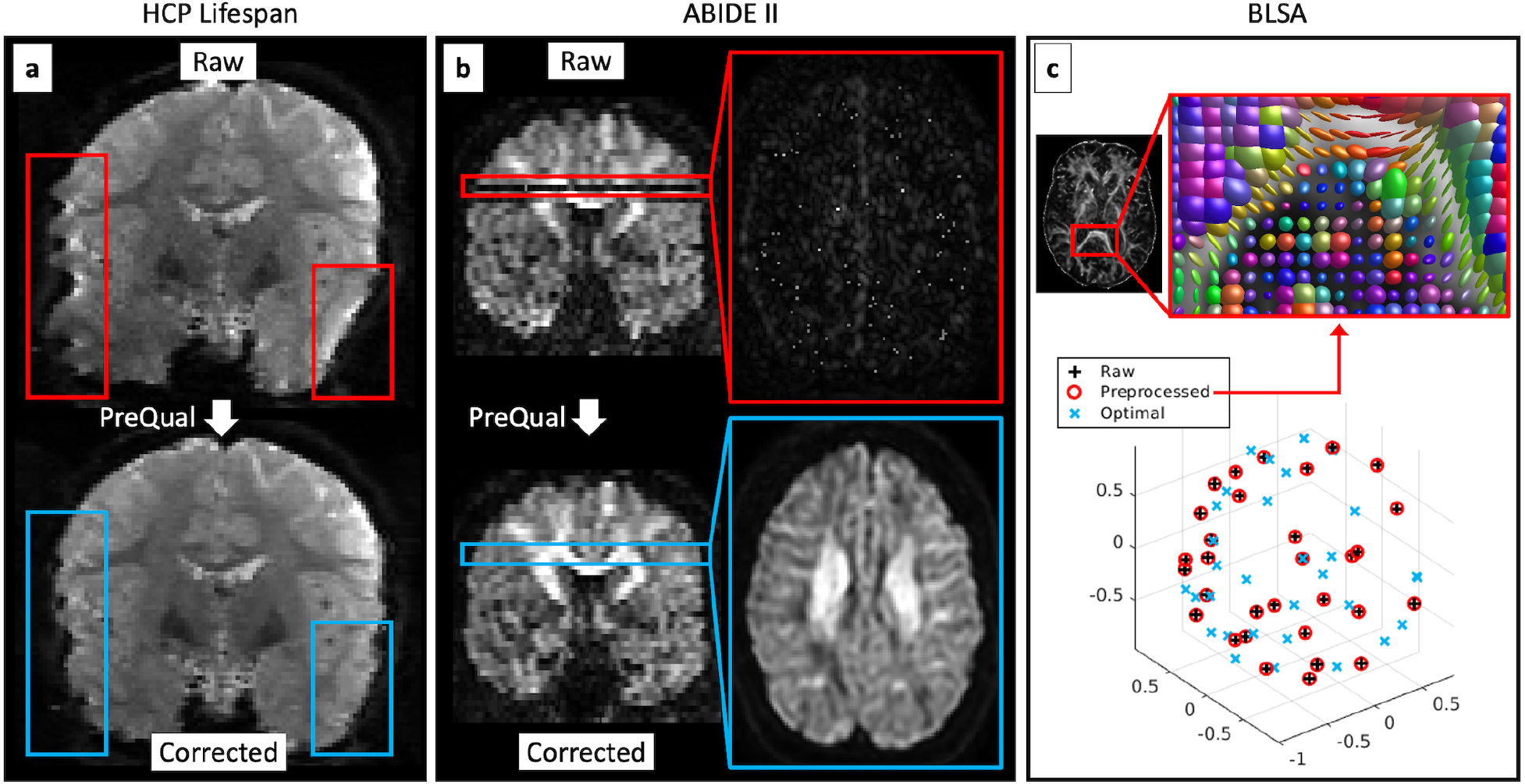
PreQual performs artifact correction and error identification on three externally available datasets. (a) For an HCP Lifespan image, PreQual corrected visually apparent susceptibility-induced distortions. The red and blue bounding boxes highlight corresponding regions before and after preprocessing by PreQual, respectively. (b) For an ABIDE II image, PreQual identified and corrected visually apparent slice-wise signal drop out. The red and blue bounding boxes highlight corresponding slices before and after preprocessing by PreQual, respectively. (c) For an uncorrected BLSA image, PreQual correctly identified improperly oriented gradients as have been previously documented and repaired via visualization of preprocessed gradients against optimal ones and visualization of tensor glyphs.

### Typical Runtimes

We quantified the runtime of PreQual on three different configurations of Dataset B (Table 1), corresponding to three forms of susceptibility-induced distortion correction: no correction, correction informed by Synb0-DisCo, and correction with traditional complementary phase encoded inputs. On a workstation running 64-bit Ubuntu 18.04 with 24 GB RAM and a 4-core 3.5 GHz Intel(R) Xeon(R) E5 processor with multithreading, we measured the runtimes to be 18 minutes and 12 seconds, 58 minutes and 47 seconds, and 48 minutes and 30 seconds, respectively. We find that under these conditions, susceptibility-induced distortion correction with FSL’s *topup* was the most time-demanding step.

**Table 1.**
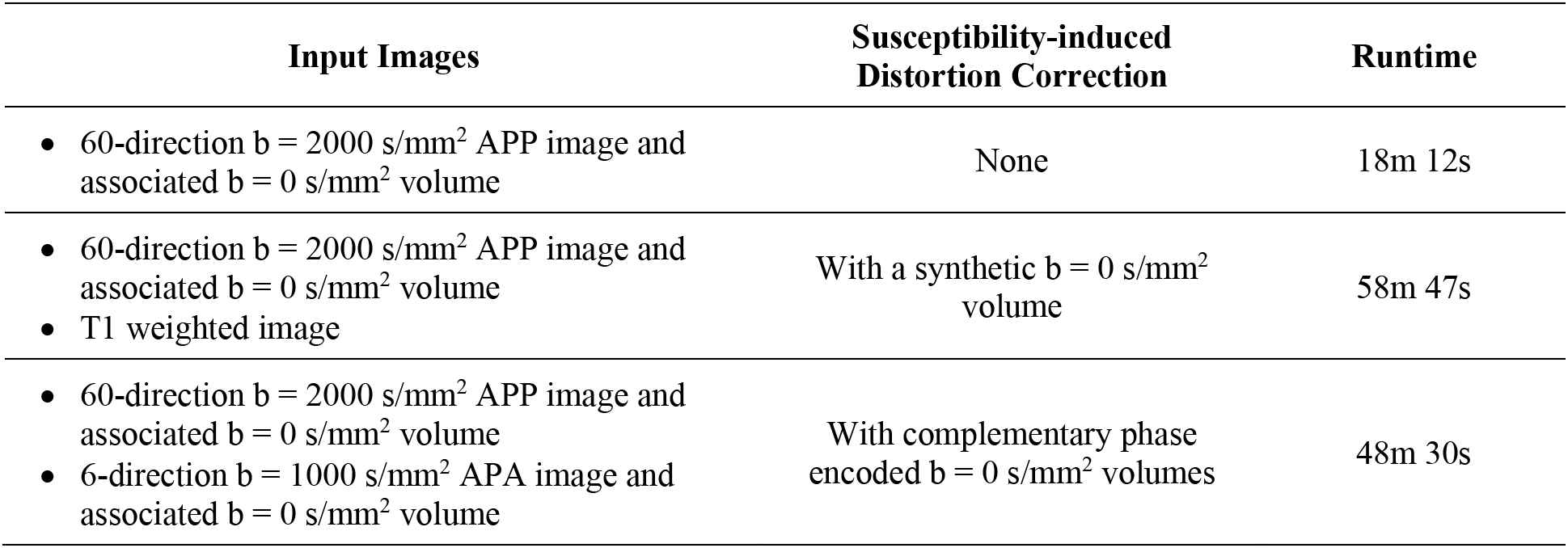
Typical runtimes of PreQual on three different configurations of Dataset B.

| Input Images | Susceptibility-induced Distortion Correction | Runtime |
| --- | --- | --- |
| 60-direction $b = 2000$ s/mm<sup>2</sup> APP image and associated $b = 0$ s/mm<sup>2</sup> volume | None | 18m 12s |
| 60-direction $b = 2000$ s/mm<sup>2</sup> APP image and associated $b = 0$ s/mm<sup>2</sup> volume<br>T1 weighted image | With a synthetic $b = 0$ s/mm<sup>2</sup> volume | 58m 47s |
| 60-direction $b = 2000$ s/mm<sup>2</sup> APP image and associated $b = 0$ s/mm<sup>2</sup> volume<br>6-direction $b = 1000$ s/mm<sup>2</sup> APA image and associated $b = 0$ s/mm<sup>2</sup> volume | With complementary phase encoded $b = 0$ s/mm<sup>2</sup> volumes | 48m 30s |

## Discussion and Conclusions

We present PreQual, a single integrated pipeline for DWI preprocessing and QA. By leveraging tools across different software packages, we are able to reproducibly apply a multitude of different preprocessing techniques to DWI data. By leveraging qualitative and quantitative QA, we demonstrate that each step of the pipeline contributes to the quality of the data and helps produce outputs better prepared for secondary modeling. We show that an analysis of gradient orientations and ROI-based average FA can identify problems with the storage and conversion of DWI data.

We envision the use of PreQual as a replacement for traditional DWI preprocessing and QA methods. Regarding preprocessing, a single pipeline reduces the need for data and software manipulation, simplifying the process and reducing the opportunity for errors and data corruption. Regarding QA, current approaches can be tedious. These include needing to open DWI images with specialized software, loop through the volumes manually, and look for artifacts and other errors; needing to hand-draw multiple ROIs for investigation; or needing to calculate and visualize model glyphs and scalar maps manually (15). Instead, we envision that all scans in a session can be run through PreQual with one command and visualization of the output QA report with a common PDF reader can provide an intuitive and efficient way for users to understand their data.

We include Gibbs de-ringing, Rician correction, and N4 bias correction and their corresponding QA steps in this pipeline (Supporting Information Figure S11). However, the authors of the Gibbs de-ringing approach have indicated that it can be unstable for partial Fourier acquisitions (55), including many EPI acquisitions with which DWI data are acquired. We also find the process to be difficult to QA: even though the artifacts often are found at high contrast boundaries, the exact boundaries and slices can be inconsistent. Further, we find Rician correction with the method of moments difficult to QA as well: although intensities are expected to decrease by the standard deviation of noise (36), we find this to be difficult to intuitively quantify. Empirically, we also find it to minimally affect results (Supporting Information Figure S13). For N4 bias field correction, we have found empirically that the fields are often orders of magnitude smaller than the intensities in the images and thus find it to be largely unnecessary. As such, we leave these features off by default and do not characterize them significantly in this study. Additionally, we note that Synb0-DisCo was trained on DWI of healthy brains, and thus we do not recommend its use on data with lesions.

A related QA tool, *eddyqc* (35), interrogates FSL’s *eddy* and provides a document that details the motion, slice-wise imputation, and other distortion-correction operations performed by FSL. With PreQual, we build on this by providing preprocessing tools from other software packages and a way to quality check other aspects of diffusion preprocessing in addition to distortion correction. We note that *eddyqc* can perform study-wise QA, something that PreQual was not explicitly designed to do. However, we save all information needed to do so in the pipeline’s outputs should the user require it (Supporting Information Figure S12). Related preprocessing tools, DESIGNER and TORTOISE, perform similar preprocessing steps with different implementations and achieve similar performance (Supporting Information Table S3 and Figure S13). These pipelines focus on preprocessing and robust modeling, and thus we complement them by focusing on preprocessing and extensive QA to prepare data for arbitrary secondary modeling.

All information presented in this manuscript was extracted directly from either pipeline outputs or from the QA documents. The pipeline was created with this design in mind in order to facilitate interrogation of the preprocessing steps for users to understand what is happening to their data. The only data reported here that are explicitly not reported in the PreQual document are the shell-wise SNRs for non-zero shells, as shown in Figure 2. Instead, in line with the *eddyqc* tool, we report the median intra-mask shell-wise CNR calculated by *eddy* in order to represent the ratio between the diffusion-induced signal variability and non-related signal variability as a measure of *eddy*’s success (35).

PreQual was designed to be modular both internally and externally. Internally, each preprocessing step aside from artifact correction and image concatenation can be turned on or off with command line options. For instance, should users choose to not perform denoising or normalization, they can do so. In these scenarios when recommended options are turned off, the output document still reports the corresponding QA metrics, allowing users to identify potential noise or intensity issues, even without performing the corresponding preprocessing steps. PreQual is also designed to take additional parameters for FSL’s *topup* and *eddy*, should users want to use more advanced features, like slice-wise intra-volume motion correction (65) or dynamic susceptibility estimation (66).

Externally, PreQual was designed to prepare DWI data for arbitrary secondary analysis. For instance, at high b-values non-Gaussian biases can affect the microstructural metrics made with tensors. In this case, a diffusion kurtosis imaging (DKI) model may be optimal for making inferences from the data. Similarly, TE dependent diffusion imaging acquisition schemes necessitate the fitting of relaxation metrics and are inadequately represented with tensors (67). In short, variation in acquisition schemes and experimental design necessarily require highly custom models on a study-by-study basis. Thus, it is not our goal to provide the best models for analysis. Instead, we intend to give all investigators the best starting point for any analysis. As such, we have selected robust preprocessing tools that work for a variety of acquisitions and a robust model, DTI, that, despite its flaws for some acquisition schemes, serves as a reproducible way of characterizing DWI data for QA purposes. In other words, although DTI may not be the best model for all DWI data, we expect that corrupt data unsuitable for secondary analysis would be identifiable with DTI. That being said, we include a warning when we detect b-values that may not be best suited for DTI modeling (i.e., less than 500 s/mm^2^ or greater than 1500 s/mm^2^) explaining that the DTI data generated by PreQual should be carefully reviewed prior to use for non-QA purposes.

Last, though primarily designed for neuroimaging applications, we anticipate that this tool will be useful for other anatomies or phantoms as well. In these cases, we anticipate the likely failure of the mask generation or white matter region identification processes. However, we provide an option to run eddy without a mask, thus allowing preprocessing and distortion correction to run on other anatomies even if some of the QA processes do not.

The PreQual source code, accompanying documentation, and a Singularity definition file for containerization have been made available to enable evaluation of the proposed pipeline at github.com/MASILab/PreQual.

## Acknowledgements

This work was conducted in part using the resources of the Advanced Computing Center for Research and Education at Vanderbilt University, Nashville, TN. This work was supported by the National Institutes of Health (NIH) under award numbers 5R01EB017230, 5T32EB001628, 5T32GM007347, 1UL1RR024975, 5R01NS110130, and 5R01NS108445. This work was also supported by the National Science Foundation under award number 1452485. This research was conducted with the support from the Intramural Research Program of the National Institute on Aging of the NIH. The content is solely the responsibility of the authors and does not necessarily represent the official views of the NIH.

## References

1. O’Donnell LJ, Westin CF. An introduction to diffusion tensor image analysis. Neurosurg. Clin. N. Am. 2011;22:185–196 doi: 10.1016/j.nec.2010.12.004.

2. Di Martino A, O’Connor D, Chen B, et al. Enhancing studies of the connectome in autism using the autism brain imaging data exchange II. Sci. Data 2017;4:170010 doi: 10.1038/sdata.2017.10.

3. Travers BG, Adluru N, Ennis C, et al. Diffusion Tensor Imaging in Autism Spectrum Disorder: A Review. Autism Res. 2012;5:289–313 doi: 10.1002/aur.1243.

4. Westlye LT, Walhovd KB, Dale AM, et al. Life-span changes of the human brain white matter: Diffusion tensor imaging (DTI) and volumetry. Cereb. Cortex 2010;20:2055–2068 doi: 10.1093/cercor/bhp280.

5. Zavaliangos-Petropulu A, Nir TM, Thomopoulos SI, et al. Diffusion MRI indices and their relation to cognitive impairment in brain aging: The updated multi-protocol approach in ADNI3. Front. Neuroinform. 2019;13:2 doi: 10.3389/fninf.2019.00002.

6. Inglese M, Bester M. Diffusion imaging in multiple sclerosis: Research and clinical implications Jensen JH, Helpern JA, editors. NMR Biomed. 2010;23:865–872 doi: 10.1002/nbm.1515.

7. De Santis S, Bastiani M, Droby A, et al. Characterizing Microstructural Tissue Properties in Multiple Sclerosis with Diffusion MRI at 7 T and 3 T: The Impact of the Experimental Design. Neuroscience 2019;403:17–26 doi: 10.1016/j.neuroscience.2018.03.048.

8. Kubicki M, McCarley R, Westin CF, et al. A review of diffusion tensor imaging studies in schizophrenia. J. Psychiatr. Res. 2007;41:15–30 doi: 10.1016/j.jpsychires.2005.05.005.

9. Cetin-Karayumak S, Di Biase MA, Chunga N, et al. White matter abnormalities across the lifespan of schizophrenia: a harmonized multi-site diffusion MRI study. Mol. Psychiatry 2019 doi: 10.1038/s41380-019-0509-y.

10. Sporns O, Tononi G, Kötter R. The human connectome: A structural description of the human brain. PLoS Comput. Biol. 2005;1:0245–0251 doi: 10.1371/journal.pcbi.0010042.

11. Bello L, Gambini A, Castellano A, et al. Motor and language DTI Fiber Tracking combined with intraoperative subcortical mapping for surgical removal of gliomas. Neuroimage 2008;39:369–382 doi: 10.1016/j.neuroimage.2007.08.031.

12. Golby AJ, Kindlmann G, Norton I, Yarmarkovich A, Pieper S, Kikinis R. Interactive diffusion tensor tractography visualization for neurosurgical planning. Neurosurgery 2011;68:496–502 doi: 10.1227/NEU.0b013e3182061ebb.

13. Raffa G, Conti A, Scibilia A, et al. The impact of diffusion tensor imaging fiber tracking of the corticospinal tract based on navigated transcranial magnetic stimulation on surgery of motor-eloquent brain lesions. Neurosurgery 2018;83:768–782 doi: 10.1093/neuros/nyx554.

14. Costabile JD, Alaswad E, D’Souza S, Thompson JA, Ormond DR. Current applications of diffusion tensor imaging and tractography in intracranial tumor resection. Front. Oncol. 2019;9:426 doi: 10.3389/fonc.2019.00426.

15. Jones DK. Diffusion MRI: theory, methods, and application. (Jones DK, editor.) Oxford; Oxford University Press; 2011.

16. Le Bihan D, Poupon C, Amadon A, Lethimonnier F. Artifacts and pitfalls in diffusion MRI. J. Magn. Reson. Imaging 2006;24:478–488 doi: 10.1002/jmri.20683.

17. Li X, Morgan PS, Ashburner J, Smith J, Rorden C. The first step for neuroimaging data analysis: DICOM to NIfTI conversion. J. Neurosci. Methods 2016;264:47–56 doi: 10.1016/j.jneumeth.2016.03.001.

18. Anderson AW, Gore JC. Analysis and correction of motion artifacts in diffusion weighted imaging. Magn. Reson. Med. 1994;32:379–387 doi: 10.1002/mrm.1910320313.

19. Jones DK, Basser PJ. “Squashing peanuts and smashing pumpkins”: How noise distorts diffusion-weighted MR data. Magn. Reson. Med. 2004;52:979–993 doi: 10.1002/mrm.20283.

20. Roalf DR, Quarmley M, Elliott MA, et al. The impact of quality assurance assessment on diffusion tensor imaging outcomes in a large-scale population-based cohort. Neuroimage 2016;125:903–919 doi: 10.1016/j.neuroimage.2015.10.068.

21. Baum GL, Roalf DR, Cook PA, et al. The impact of in-scanner head motion on structural connectivity derived from diffusion MRI. Neuroimage 2018;173:275–286 doi: 10.1016/j.neuroimage.2018.02.041.

22. Oldham S, ArnatkeviČiūtė A, Smith RE, Tiego J, Bellgrove MA, Fornito A. The efficacy of different preprocessing steps in reducing motion-related confounds in diffusion MRI connectomics. Neuroimage 2020;222 doi: 10.1016/j.neuroimage.2020.117252.

23. Yendiki A, Koldewyn K, Kakunoori S, Kanwisher N, Fischl B. Spurious group differences due to head motion in a diffusion MRI study. Neuroimage 2014;88:79–90 doi: 10.1016/j.neuroimage.2013.11.027.

24. Veraart J, Novikov DS, Christiaens D, Ades-aron B, Sijbers J, Fieremans E. Denoising of diffusion MRI using random matrix theory. Neuroimage 2016;142:394–406 doi: 10.1016/j.neuroimage.2016.08.016.

25. Veraart J, Fieremans E, Novikov DS. Diffusion MRI noise mapping using random matrix theory. Magn. Reson. Med. 2016;76:1582–1593 doi: 10.1002/mrm.26059.

26. Cordero-Grande L, Christiaens D, Hutter J, Price AN, Hajnal J V. Complex diffusion-weighted image estimation via matrix recovery under general noise models. Neuroimage 2019;200:391–404 doi: 10.1016/j.neuroimage.2019.06.039.

27. Andersson JLR, Skare S, Ashburner J. How to correct susceptibility distortions in spin-echo echo-planar images: Application to diffusion tensor imaging. Neuroimage 2003;20:870–888 doi: 10.1016/S1053-8119(03)00336-7.

28. Smith SM, Jenkinson M, Woolrich MW, et al. Advances in functional and structural MR image analysis and implementation as FSL. Neuroimage 2004;23:S208–S219 doi: 10.1016/j.neuroimage.2004.07.051.

29. Schilling KG, Blaber J, Hansen C, et al. Distortion correction of diffusion weighted MRI without reverse phase-encoding scans or field-maps. PLoS One 2020;15:e0236418.

30. Andersson JLR, Sotiropoulos SN. An integrated approach to correction for off-resonance effects and subject movement in diffusion MR imaging. Neuroimage 2016;125:1063–1078 doi: 10.1016/j.neuroimage.2015.10.019.

31. Andersson JLR, Graham MS, Zsoldos E, Sotiropoulos SN. Incorporating outlier detection and replacement into a non-parametric framework for movement and distortion correction of diffusion MR images. Neuroimage 2016;141:556–572 doi: 10.1016/j.neuroimage.2016.06.058.

32. Jeurissen B, Leemans A, Sijbers J. Automated correction of improperly rotated diffusion gradient orientations in diffusion weighted MRI. Med. Image Anal. 2014;18:953–962 doi: 10.1016/j.media.2014.05.012.

33. Lauzon CB, Asman AJ, Esparza ML, et al. Simultaneous Analysis and Quality Assurance for Diffusion Tensor Imaging Alexander DC, editor. PLoS One 2013;8:e61737 doi: 10.1371/journal.pone.0061737.

34. Oguz I, Farzinfar M, Matsui J, et al. DTIPrep: Quality control of diffusion-weighted images. Front. Neuroinform. 2014;8:4 doi: 10.3389/fninf.2014.00004.

35. Bastiani M, Cottaar M, Fitzgibbon SP, et al. Automated quality control for within and between studies diffusion MRI data using a non-parametric framework for movement and distortion correction. Neuroimage 2019;184:801–812 doi: 10.1016/j.neuroimage.2018.09.073.

36. Ades-Aron B, Veraart J, Kochunov P, et al. Evaluation of the accuracy and precision of the diffusion parameter EStImation with Gibbs and NoisE removal pipeline. Neuroimage 2018;183:532–543 doi: 10.1016/j.neuroimage.2018.07.066.

37. Pierpaoli C, Walker L. TORTOISE: an integrated software package for processing of diffusion MRI data. 18th Annu. Meet. Int. Soc. Magn. Reson. Med. 2010;51:2010.

38. Irfanoglu MO, Nayak A, Jenkins J, Pierpaoli C. TORTOISE v3: Improvements and New Features of the NIH Diffusion MRI Processing Pipeline. In: 25th Annual Meeting of the International Society fro Magnetic Resonance in Medicine.; 2017. pp. 9–11. doi: 10.1016/j.neuroimage.2016.08.016.5.

39. Irfanoglu MO, Modi P, Nayak A, Hutchinson EB, Sarlls J, Pierpaoli C. DR-BUDDI (Diffeomorphic Registration for Blip-Up blip-Down Diffusion Imaging) method for correcting echo planar imaging distortions. Neuroimage 2015;106:284–299 doi: 10.1016/j.neuroimage.2014.11.042.

40. Cieslak M, Cook PA, He X, et al. QSIPrep: An integrative platform for preprocessing and reconstructing diffusion MRI. bioRxiv 2020:2020.09.04.282269.

41. Tournier JD, Smith R, Raffelt D, et al. MRtrix3: A fast, flexible and open software framework for medical image processing and visualisation. Neuroimage 2019;202:116137 doi: 10.1016/j.neuroimage.2019.116137.

42. Jenkinson M, Beckmann CF, Behrens TEJ, Woolrich MW, Smith SM. FSL. Neuroimage 2012;62:782–790 doi: 10.1016/j.neuroimage.2011.09.015.

43. Tustison NJ, Cook PA, Klein A, et al. Large-scale evaluation of ANTs and FreeSurfer cortical thickness measurements. Neuroimage 2014;99:166–179 doi: 10.1016/j.neuroimage.2014.05.044.

44. Smith SM. Fast robust automated brain extraction. Hum. Brain Mapp. 2002;17:143–155 doi: 10.1002/hbm.10062.

45. Nelder JA, Mead R. A Simplex Method for Function Minimization. Comput. J. 1965;7:308–313 doi: 10.1093/comjnl/7.4.308.

46. Veraart J, Sijbers J, Sunaert S, Leemans A, Jeurissen B. Weighted linear least squares estimation of diffusion MRI parameters: Strengths, limitations, and pitfalls. Neuroimage 2013;81:335–346 doi: 10.1016/j.neuroimage.2013.05.028.

47. Westin CF, Peled S, Gudbjartsson H, Kikinis R, Jolesz FA. Geometrical Diffusion Measures for MRI from Tensor Basis Analysis. Proc. 5th Annu. Meet. ISMRM 1997:1742.

48. Tournier JD, Calamante F, Connelly A. MRtrix: Diffusion tractography in crossing fiber regions Lee J, editor. Int. J. Imaging Syst. Technol. 2012;22:53–66 doi: 10.1002/ima.22005.

49. Mori S, Wakana S, Van Zijl PCM, Nagae-Poetscher LM. MRI atlas of human white matter. Elsevier; 2005.

50. Wakana S, Caprihan A, Panzenboeck MM, et al. Reproducibility of quantitative tractography methods applied to cerebral white matter. Neuroimage 2007;36:630–644 doi: 10.1016/j.neuroimage.2007.02.049.

51. Hua K, Zhang J, Wakana S, et al. Tract probability maps in stereotaxic spaces: Analyses of white matter anatomy and tract-specific quantification. Neuroimage 2008;39:336–347 doi: 10.1016/j.neuroimage.2007.07.053.

52. Avants BB, Epstein CL, Grossman M, Gee JC. Symmetric diffeomorphic image registration with cross-correlation: Evaluating automated labeling of elderly and neurodegenerative brain. Med. Image Anal. 2008;12:26–41 doi: 10.1016/j.media.2007.06.004.

53. Kochunov P, Glahn DC, Lancaster J, et al. Fractional anisotropy of cerebral white matter and thickness of cortical gray matter across the lifespan. Neuroimage 2011;58:41–49 doi: 10.1016/j.neuroimage.2011.05.050.

54. Zhang Y, Brady M, Smith S. Segmentation of brain MR images through a hidden Markov random field model and the expectation-maximization algorithm. IEEE Trans. Med. Imaging 2001;20:45–57 doi: 10.1109/42.906424.

55. Kellner E, Dhital B, Kiselev VG, Reisert M. Gibbs-ringing artifact removal based on local subvoxel-shifts. Magn. Reson. Med. 2016;76:1574–1581 doi: 10.1002/mrm.26054.

56. Koay CG, Basser PJ. Analytically exact correction scheme for signal extraction from noisy magnitude MR signals. J. Magn. Reson. 2006;179:317–322 doi: 10.1016/j.jmr.2006.01.016.

57. Tustison NJ, Avants BB, Cook PA, et al. N4ITK: Improved N3 bias correction. IEEE Trans. Med. Imaging 2010;29:1310–1320 doi: 10.1109/TMI.2010.2046908.

58. Gudbjartsson H, Patz S. The Rician distribution of noisy MRI data. Magn. Reson. Med. 1995;34:910–914 doi: 10.1002/mrm.1910340618.

59. Cai LY, Yang Q, Kanakaraj P, et al. MASiVar: Multisite, Multiscanner, and Multisubject Acquisitions for Studying Variability in Diffusion Weighted Magnetic Resonance Imaging. bioRxiv 2020:2020.12.03.408567 doi: 10.1101/2020.12.03.408567.

60. Harms MP, Somerville LH, Ances BM, et al. Extending the Human Connectome Project across ages: Imaging protocols for the Lifespan Development and Aging projects. Neuroimage 2018;183:972–984 doi: 10.1016/j.neuroimage.2018.09.060.

61. Shock NW. Normal human aging: the Baltimore longitudinal study of aging. (Shock NW (Nathan W, editor.) Baltimore, Md: U.S. Dept. of Health and Human Services, Public Health Service, National Institutes of Health, National Institute on Aging, Gerontology Research Center; 1984.

62. Resnick SM, Goldszal AF, Davatzikos C, et al. One-year age changes in MRI brain volumes in older adults. Cereb. Cortex 2000;10:464–472 doi: 10.1093/cercor/10.5.464.

63. Schilling KG, Yeh FC, Nath V, et al. A fiber coherence index for quality control of B-table orientation in diffusion MRI scans. Magn. Reson. Imaging 2019;58:82–89 doi: 10.1016/j.mri.2019.01.018.

64. Hollander M, Wolfe DA, Chicken E. Nonparametric statistical methods. John Wiley & Sons; 2013.

65. Andersson JLR, Graham MS, Drobnjak I, Zhang H, Filippini N, Bastiani M. Towards a comprehensive framework for movement and distortion correction of diffusion MR images: Within volume movement. Neuroimage 2017;152:450–466 doi: 10.1016/j.neuroimage.2017.02.085.

66. Andersson JLR, Graham MS, Drobnjak I, Zhang H, Campbell J. Susceptibility-induced distortion that varies due to motion: Correction in diffusion MR without acquiring additional data. Neuroimage 2018;171:277–295 doi: 10.1016/j.neuroimage.2017.12.040.

67. Veraart J, Novikov DS, Fieremans E. TE dependent Diffusion Imaging (TEdDI) distinguishes between compartmental T2 relaxation times. NeuroImage (Orlando, Fla.) 182:360–369 doi: 10.1016/j.neuroimage.2017.09.030.

68. Dhollander T, Raffelt D, Connelly A. Unsupervised 3-tissue response function estimation from single-shell or multi-shell diffusion MR data without a co-registered T1 image. ISMRM Work. Break. Barriers Diffus. MRI 2016:5.

69. Iglesias JE, Liu CY, Thompson PM, Tu Z. Robust brain extraction across datasets and comparison with publicly available methods. IEEE Trans. Med. Imaging 2011;30:1617–1634 doi: 10.1109/TMI.2011.2138152.

70. Nobuyuki Otsu. A Threshold Selection Method from Gray-Level Histograms. IEEE Trans. Syst. Man Cybern 1979;9:62–66.

