## Supporting Information for "PreQual: An automated pipeline for integrated preprocessing and quality assurance of diffusion weighted MRI images"

We supply Supporting Information Table S1 explaining characteristics of successful DICOM to NIFTI conversions, signs of conversion failure, and first steps to resolve them.

**Supporting Information Table S1.** DICOM to NIFTI conversions.

| Characteristics of successful conversions | Hallmarks of conversion failure… | …and how to begin to fix them |
|---|---|---|
| Gradients are re-oriented to subject space from scanner space | • The vectors in the .bvec file and in the DICOM header match exactly and the Image Orientation Patient (IOP) field in the DICOM header is not [1, 0, 0, 0, 1, 0].<br>• Tractography for robust white matter bundles inexplicably fails.<br>• Tensors are rotated away from expected physiological orientations. | • Use a recent version of *dcm2niix*.<br>• Manually re-orient the gradients. The first three and last three elements in the IOP field reflect the three direction cosines of the first row and first column with respect to the patient. The rotation matrix reorienting the b-vectors to subject space can be constructed as $R = \begin{bmatrix} x^T \\ y^T \\ z^T \end{bmatrix}$, where $x = \begin{bmatrix} IOP(1) \\ -IOP(2) \\ IOP(3) \end{bmatrix}$, $y = \begin{bmatrix} -IOP(4) \\ IOP(5) \\ -IOP(6) \end{bmatrix}$, and $z = x \times y$ |
| The NIFTI volumes are in the order they were acquired on the scanner | • The FA analysis produces unrealistic regional values and the FA maps do not appear as expected.<br>• The b = 0 s/mm$^2$ volume was set to be acquired last on the exam card but appears first in the NIFTI sequence. | • Use a recent version of *dcm2niix*. |

We supply Supporting Information Figures S1 through S10 displaying the individual pages of the pipeline QA document, language detailing how to best interpret them, and how to identify and fix errors. They were generated on one session from each of three externally available datasets (Supporting Information Table S2). These sessions were previously identified to have isolated QA issues (29,63).

**Supporting Information Table S2.** Externally available datasets. These three sessions were processed with PreQual. The pages of the output QA documents are presented in Supporting Information Figures S1 through S10.

| **Project** | HCP Lifespan Cohort | ABIDE II | BLSA |
|---|---|---|---|
| **Site** | N/A | Barrow Neurological Institute | N/A |
| **Subject** | 2001 | 29028 | 1044 |
| **Session** | 1 | 1 | 18-0_07 |
| **Scan(s)** | • 75-direction multi-shell b = 1000 and 2500 s/mm$^2$ right to left phase encoded (RL) with 5 interspersed b = 0 s/mm$^2$ volumes<br>• 75-direction multi-shell b = 1000 and 2500 s/mm$^2$ left to right phase encoded (LR) with 5 interspersed b = 0 s/mm$^2$ volumes<br>• 76-direction multi-shell b = 1000 and 2500 s/mm$^2$ RL with 5 interspersed b = 0 s/mm$^2$ volumes<br>• 76-direction multi-shell b = 1000 and 2500 s/mm$^2$ LR with 5 interspersed b = 0 s/mm$^2$ volumes | • 32-direction b = 2500 s/mm$^2$ APP with 1 b = 0 s/mm$^2$ volume<br>• T1 anatomical image | • 30-direction b = 700 s/mm$^2$ APP with 1 b = 0 s/mm$^2$ volume<br>• T1 anatomical image |

We supply Supporting Information Figures S11 and S12 to detail additional pipeline options and considerations. We provide Supporting Information Table S3 and Figure S13 to compare PreQual to the DESIGNER and TORTOISE pipelines.

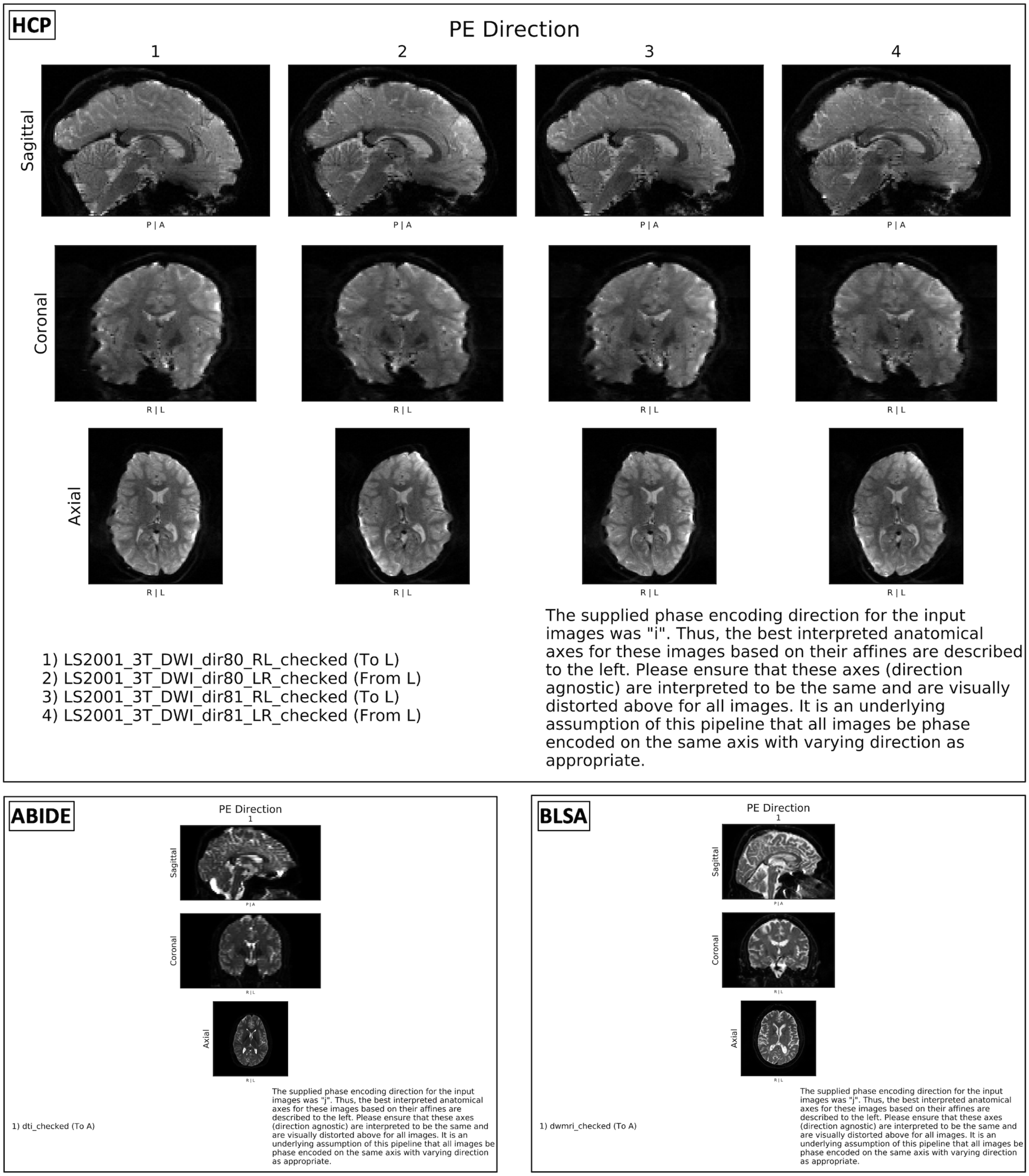


**Supporting Information Figure S1.** QA of the input images and phase encoding schemes. The report visualizes the first b = 0 s/mm$^2$ volume of each raw input image and their corresponding input phase encoding schemes with anatomical context for user verification. When interpreting these figures, we verify that the susceptibility-induced distortions are along the named axis. We also verify that all images with a shared direction along the axis exhibit similar morphology distinct from scans acquired in the other direction. For instance, in the HCP Lifespan session, scans 1 and 3 are encoded from right-to-left and scans 2 and 4 from left-to-right. We confirm that the first pair of scans are "smeared" in a similar direction that is opposite to the "smear" of the images in the second pair. We note that with *topup*, it does not matter if the right-to-left scans are actually right-to-left phase encoded, as long as they are all the same direction and opposite to those specified as left-to-right. This generalizes to the posterior-to-anterior axis as well. If we identify errors or inconsistencies, we suggest first ensuring that the directions specified in the configuration file are accurate. We then suggest reviewing the DICOM headers and/or the exam protocols.

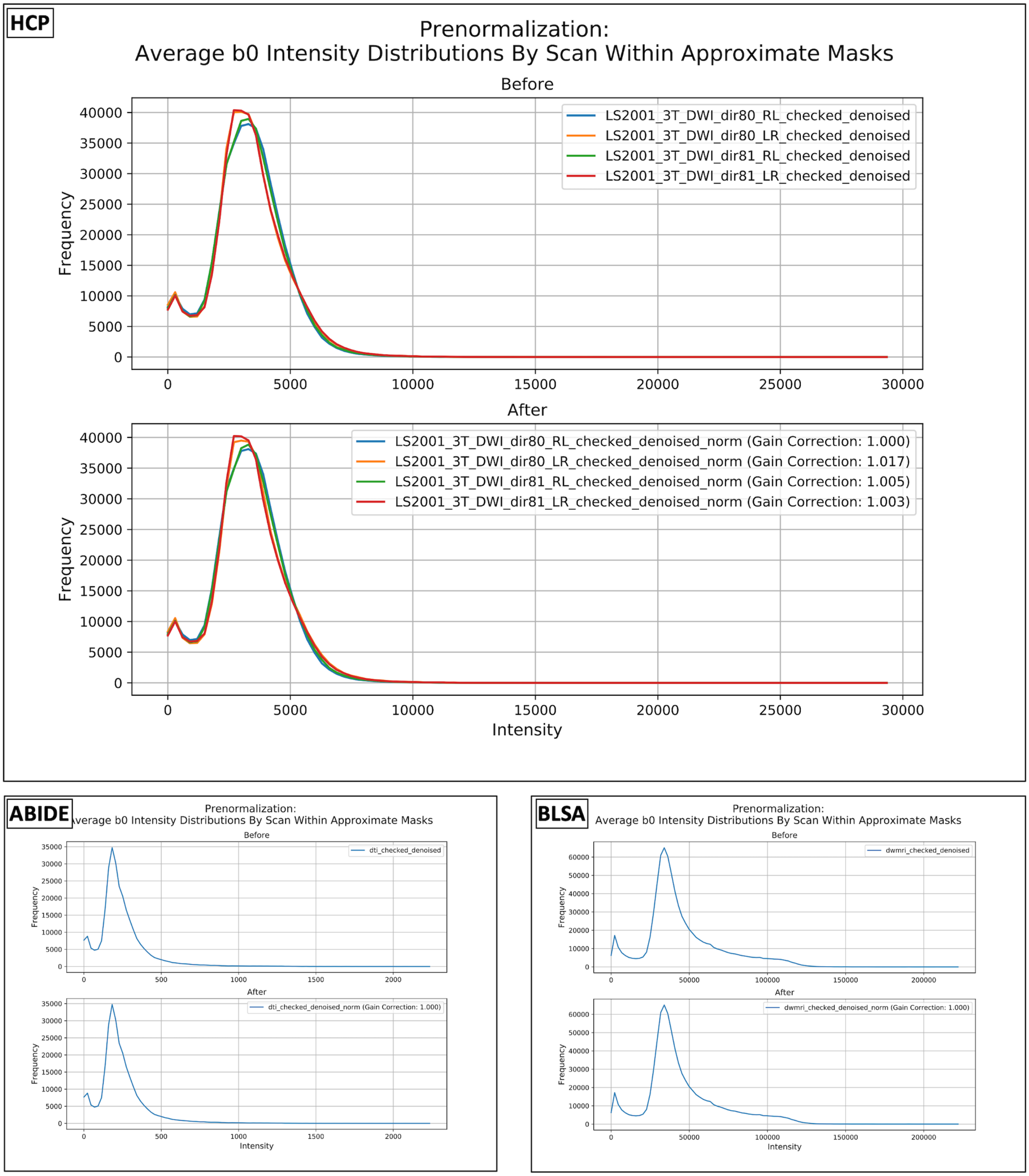


**Supporting Information Figure S2.** QA of the normalization step. The report details the intensity histograms of the average b = 0 s/mm$^2$ volume from each input image before and after normalization and reports the calculated scale factors that maximize their intensity histogram intersections. When interpreting these figures, we seek to identify any sessions with potential prescan gain change effects and confirm they were corrected. This is identified via the calculation of correction factors roughly more than 1.05 and less than 0.95. If this is the case, we verify the distributions after normalization overlap and that the median intensities of the different shells exhibit expected physiologic relationships (Supporting Information Figure S5) as an indication of successful normalization. If it was not successful, we first investigate the affected images for any obvious intensity-based artifacts. In the case of the HCP Lifespan session, likely no gain changes were present since the calculated scale factors were all close to 1. At this point we could consider rerunning preprocessing without normalization as to not induce artificial intensity changes. Empirically, however, we do not find that normalization of sessions without prescan impacts data quality significantly. In the case of the ABIDE II and BLSA sessions, only one input image was supplied and thus no normalization was performed.

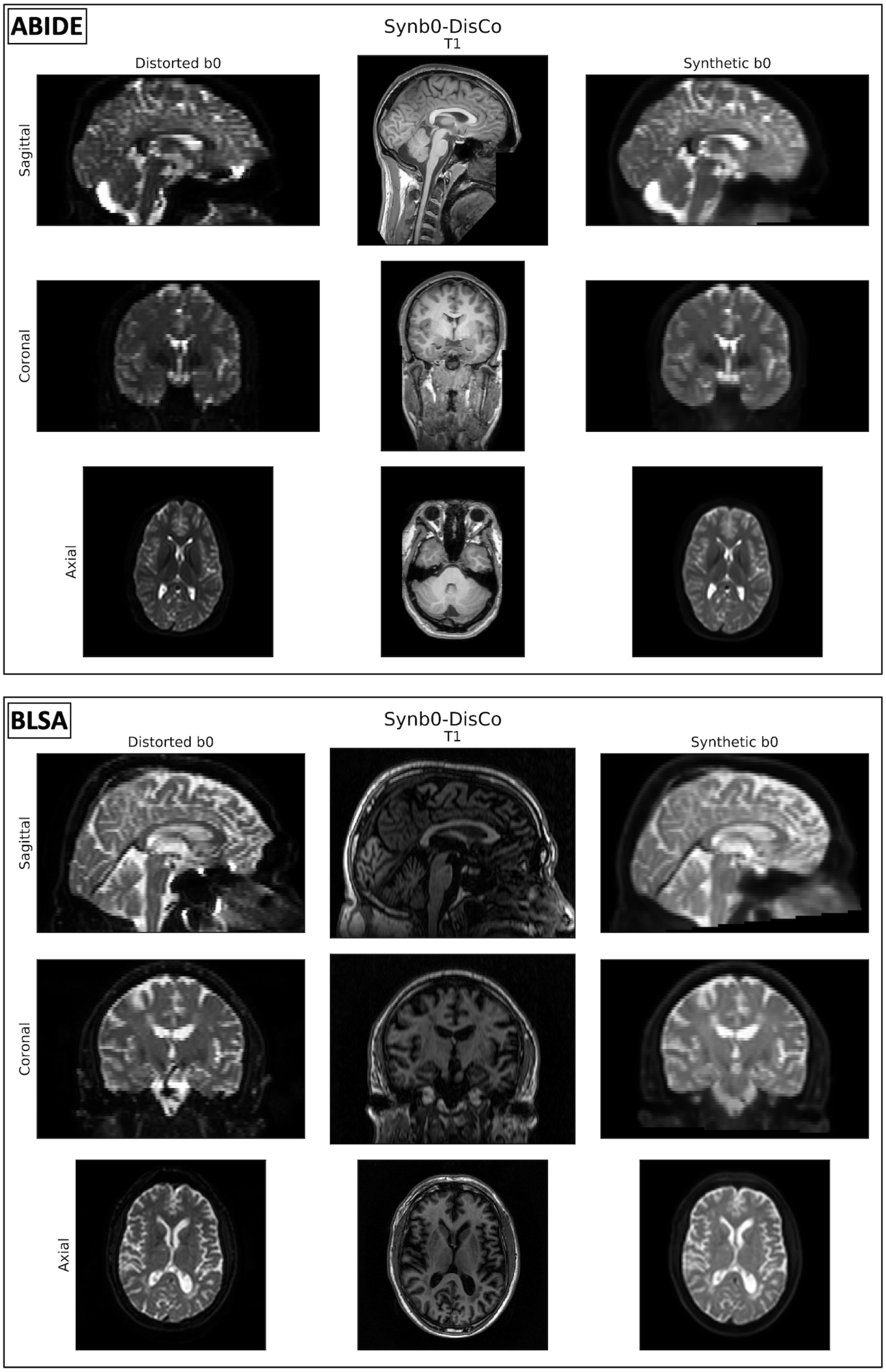


**Supporting Information Figure S3.** QA of Synb0-DisCo. The report displays the distorted b = 0 s/mm$^2$ (the first of the first input image) and T1 volume input into Synb0-DisCo and the synthetic output. The synthetic output must be used in conjunction with FSL's *topup* to perform susceptibility-induced distortion correction on the remainder of the input images. HCP Lifespan did not require Synb0-DisCo, as complementary phase encoded images were supplied. We use these figures to understand the success of Synb0-DisCo. We verify that the synthetic b = 0 s/mm$^2$ volume has similar contrast to the distorted b = 0 s/mm$^2$ volume but exhibits consistent morphology with the T1 weighted image. As a secondary check, we refer to the "Preprocessing and Masks" page (Supporting Information Figure S4) to see the success of susceptibility-induced distortion correction with *topup* and the synthetic volume by looking at the morphology changes between the raw images and the preprocessed one. If errors in Synb0-DisCo are identified, we refer to the Synb0-DisCo documentation at github.com/MASILab/Synb0-DisCo.

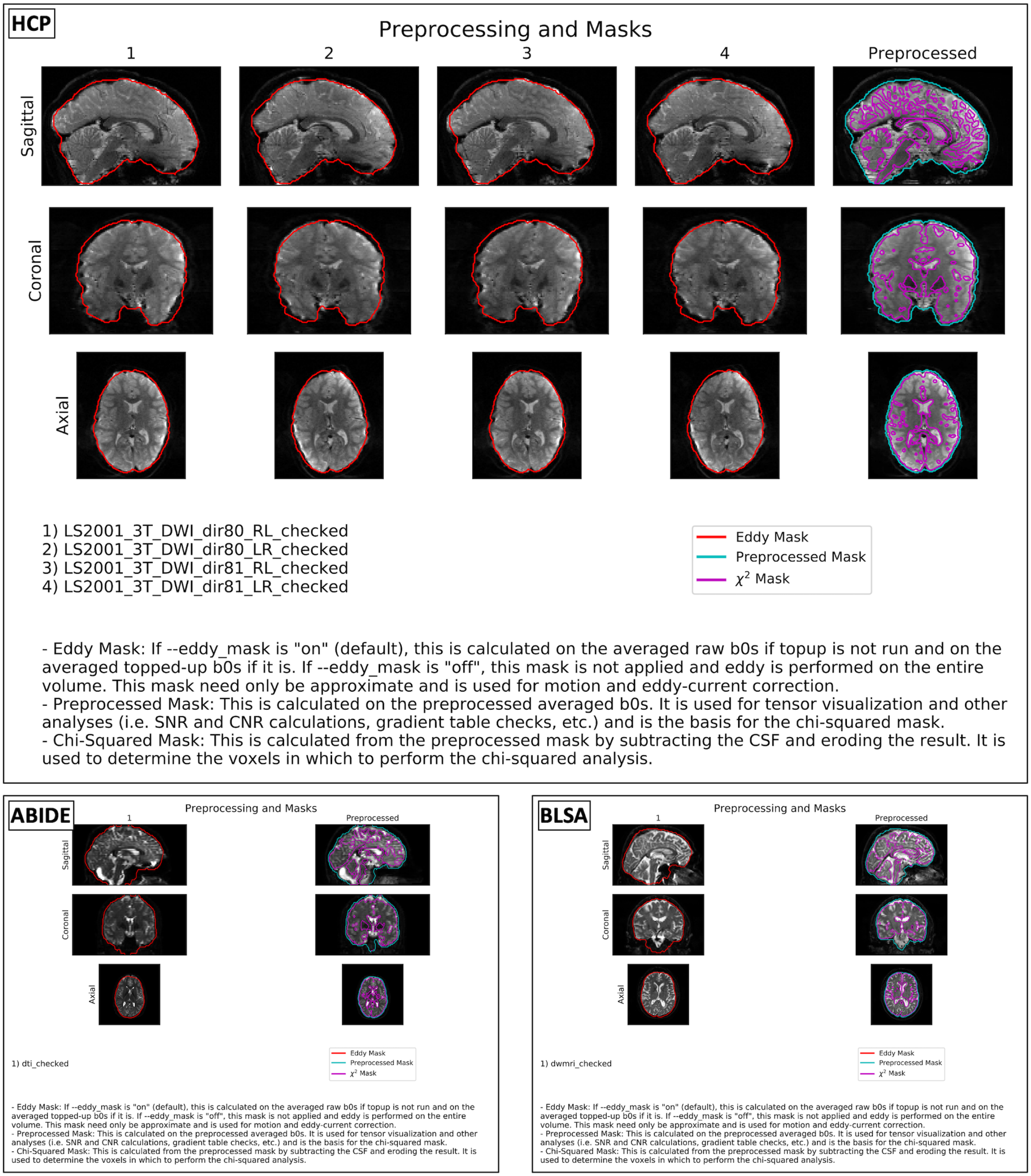


**Supporting Information Figure S4.** QA of masks. The report shows the first b = 0 s/mm$^2$ volume of each input image as well as that of the preprocessed output image. Contours of the three primary masks used in the pipeline are overlaid: the mask for *eddy*, the final mask calculated on preprocessed outputs, and the parenchyma mask used for the chi-squared goodness-of-fit analysis. When interpreting these figures, we first verify the quality of the masking process. We ensure that the mask for *eddy* and the preprocessed mask do not omit large regions of the brain and that the chi-squared mask largely excludes CSF and is restricted to brain parenchyma. If we identify errors in the *eddy* mask, we consider rerunning the pipeline without a mask at all. If we identify errors in the chi-squared mask, we make a note when interpreting the goodness-of-fit plots (Supporting Information Figure S5), as poor masking can corrupt this analysis. If we identify errors in the preprocessing mask, we consider generating an alternative mask prior to secondary analysis. Errors here are often due to FSL's *bet* failure, thus we recommend either rerunning *bet* externally with different parameters or pursuing alternative approaches such as the *dwi2mask* tool from MRTrix3 (68), the *ROBEX* tool if a T1 is available (69), or the *median_otsu* method implemented in DIPY (70). We also use this figure as a heuristic for distortion correction success. By comparing the distorted regions in the raw volumes to those in the preprocessed volume, we get a feel for how well the distortions were corrected. In the HCP Lifespan and BLSA sessions, we see markedly improved distortions.

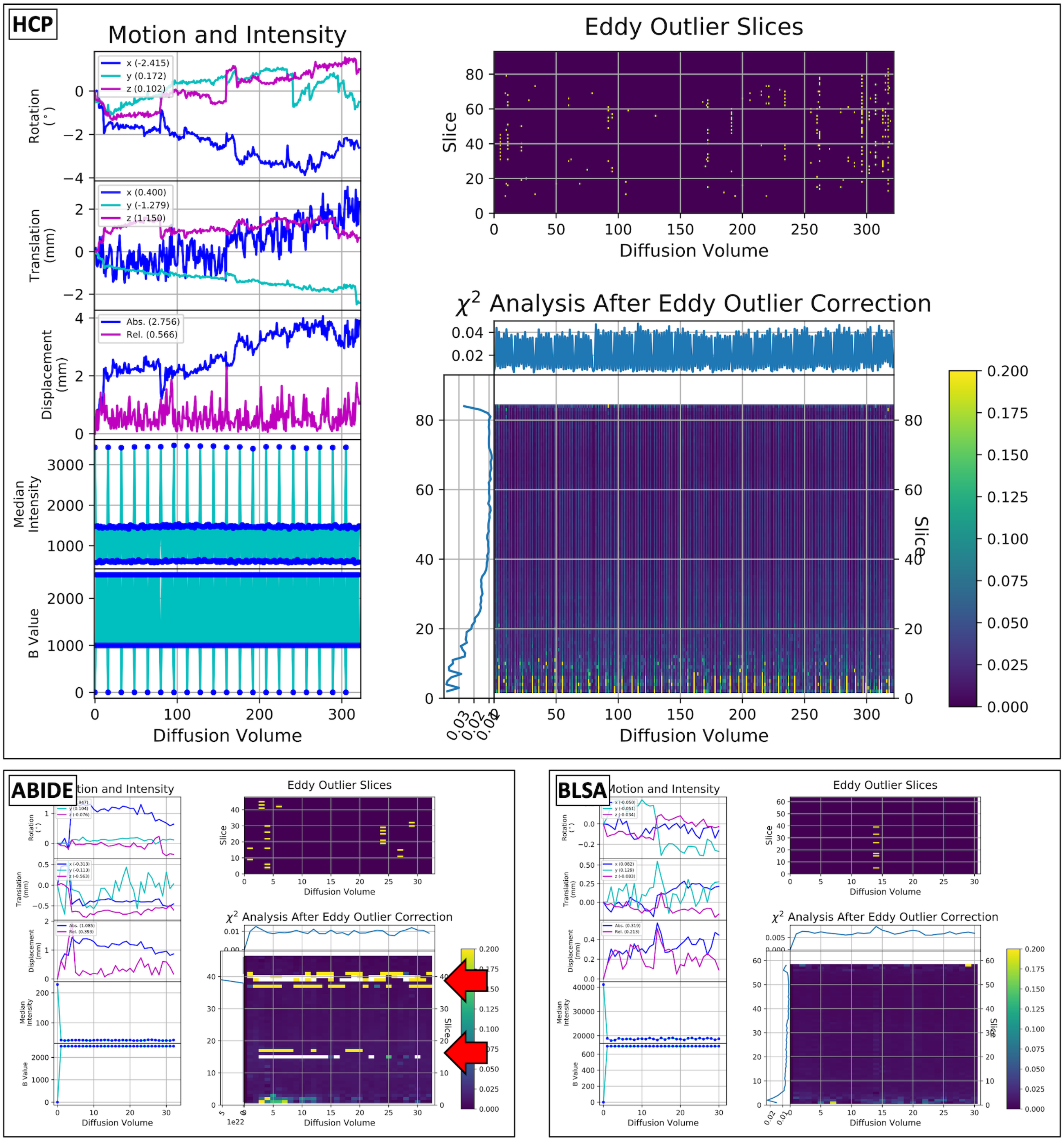


**Supporting Information Figure S5.** QA of motion correction, slice-wise signal dropout imputation, and chi-squared analysis. The inter-volume rotation, translation, and displacement; median intra-mask intensity; and b-values of the preprocessed outputs are plotted on the left side. Imputed slices are indicated on the top right of each page. The per-slice per-volume chi-squared values and their median profiles in each dimension are displayed on the bottom right of each page. The chi-squared plot color-axis is bound from 0 to 0.2 as detailed by Lauzon et al. (33). When interpreting these plots, we first look at the motion plots. We confirm that the average summary statistics listed in the legends are representative of the plots and that the magnitudes of motion are not unrealistic. If they are unrealistic, we further investigate by rendering the raw volumes. Next, we look at the median intensity and b-value plots to ensure that the intensities of the different shells have expected physical relationships: intensity should decrease with shell size and intensities within the same shell should be roughly equal. This process should be used in conjunction with interpretation of the normalization process (Supporting Information Figure S2). We then look at the outlier plot to ensure that the outlier identification process employed by *eddy* did not identify more outliers than not. If we identify a concerning plot, we first visualize the volumes. If it appears there are fewer dropped slices than identified, we consider rerunning the pipeline and modulating the outlier threshold using the extra *eddy* option *–ol_nstd*. Otherwise, we consider omitting the dataset from further analysis. Last, we interpret the chi-squared goodness-of-fit plot. As described by Lauzon et al, we expect to see values hovering just above 0 and values near or above 0.2 (33). We expect to see the high values near the minimum and maximum slices, as there is little brain parenchyma for fitting there. Similarly, in the ABIDE II session, the white and yellow values in the chi-squared plot (red arrows) across many volumes around slices 15 and 40 are due to poor masking of the brain parenchyma. Incorporation of CSF, background, and poorly fit anatomy within the middle slices of volumes often results in either unrealistically high (yellow) or infinite (white) chi-squared values. Slices with no brain parenchyma are "Not a Number" (white) across all volumes. We expect good fits to have the majority of the plots be non-yellow and non-white, as demonstrated in these three fits. If this is not the case, we suggest visualization of the tensor glyphs as a first step in understanding the fitting process (Supporting Information Figure S8).

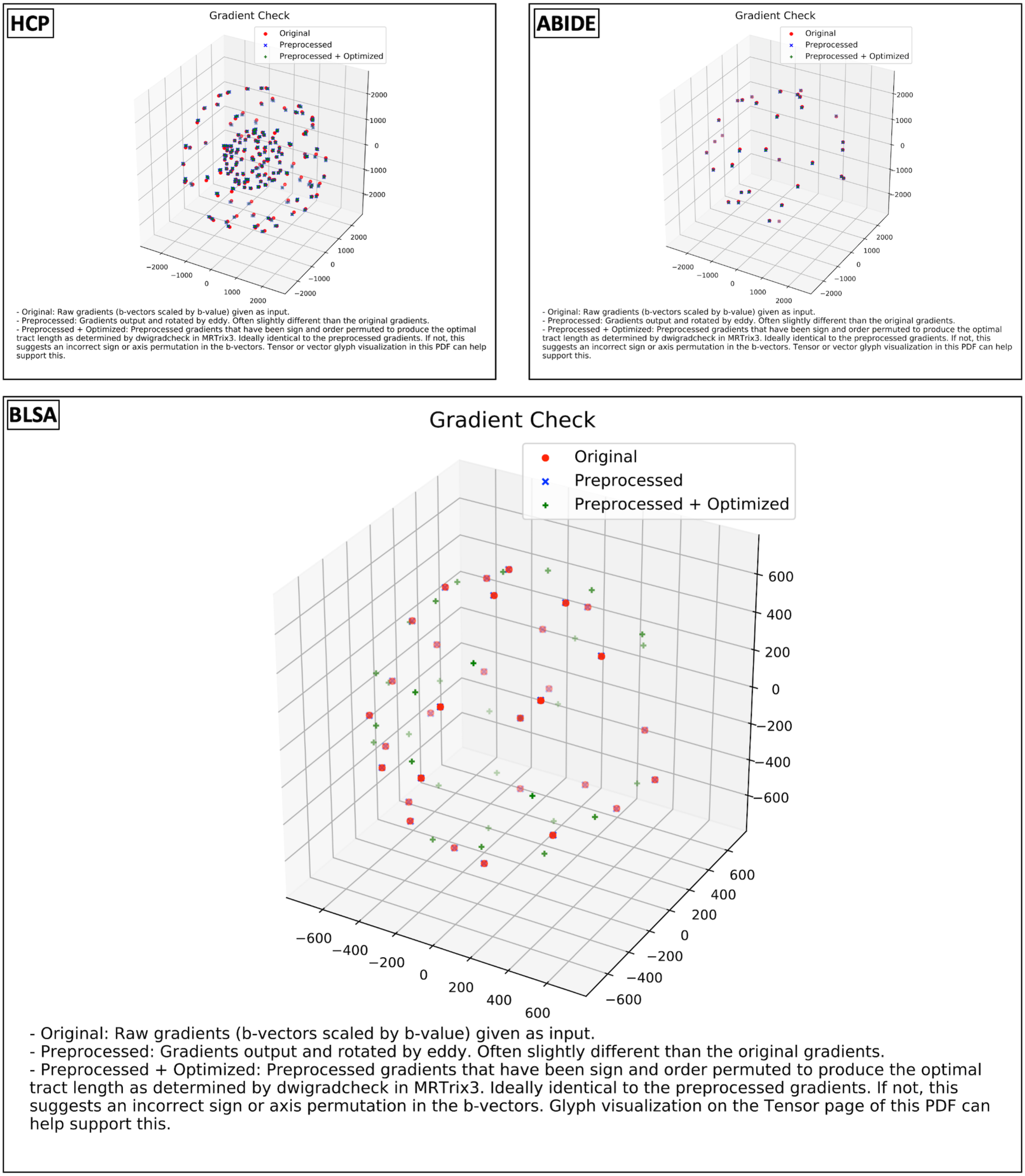


**Supporting Information Figure S6.** QA of gradients. The report plots the original input gradients, those output by eddy (preprocessed), and those determined to be sign and axis optimal by MRTrix3's *dwigradcheck*. Each point represents a b-vector scaled by its corresponding b-value. When interpreting these plots, we first confirm that differences between the preprocessed and optimal gradients do not exist, indicating that the gradients are oriented properly. Because this process depends on tractography and is intended to QA the raw gradients, we use the preprocessed gradients as a surrogate. Thus, we then confirm that only minor rotations between the original and preprocessed gradients exist, thus verifying the validity of this approach. If we identify inconsistencies with either of these expectations, as is shown in the BLSA session, we first verify that the resultant tensors are oriented improbably (Supporting Information Figure S8). Because gradients are often not simply affected by sign or axis errors, we do not overwrite the preprocessed gradients with the optimal ones identified by this analysis. Rather, we suggest users investigate potential issues impacting the gradients. First, we verify that the gradients are in the same coordinate space and have the same sign expectations as the FSL format. Notably, MRTrix3 flips the first axis. We then investigate potential reorientation errors during DICOM to NIFTI conversion (Supporting Information Table S1).

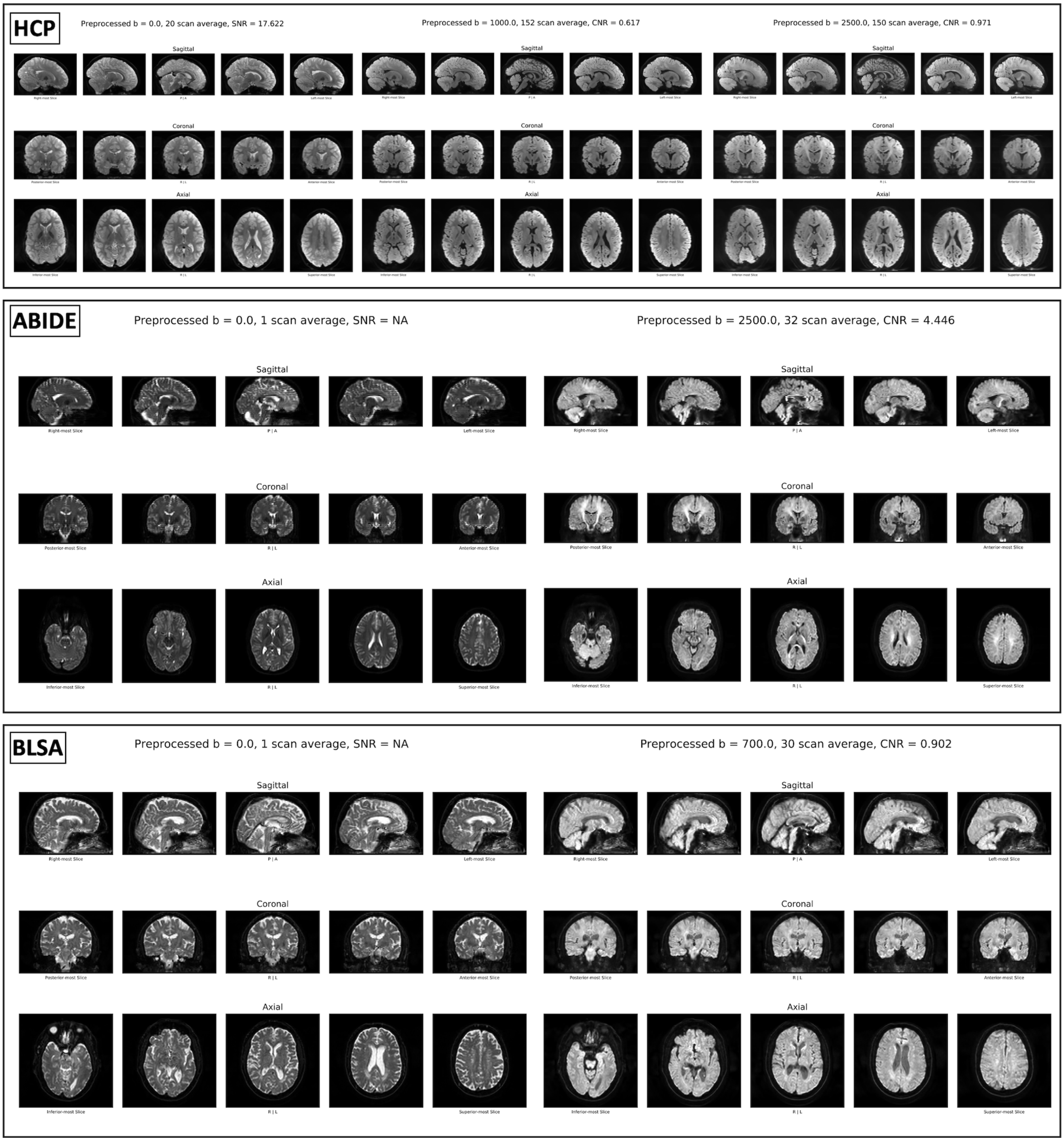


**Supporting Information Figure S7.** QA of preprocessed shells and denoising. The report visualizes five central triplanar slices of the shell-wise average volumes alongside the median intra-mask SNR and CNR. The SNR is reported as “Not a Number” when only one b = 0 s/mm$^2$ volume exists, as is the case for the ABIDE II and BLSA sessions. When interpreting these plots, we first verify that the volumes appear undistorted. If they do not, we investigate the phase-encoding configurations passed into the pipeline (Supporting Information Figure S1). We then verify the SNR and CNR. We expect the SNR for the b = 0 s/mm$^2$ volumes to be roughly 10 to 30. We expect the CNR measurements to be roughly 0.5 to 5.

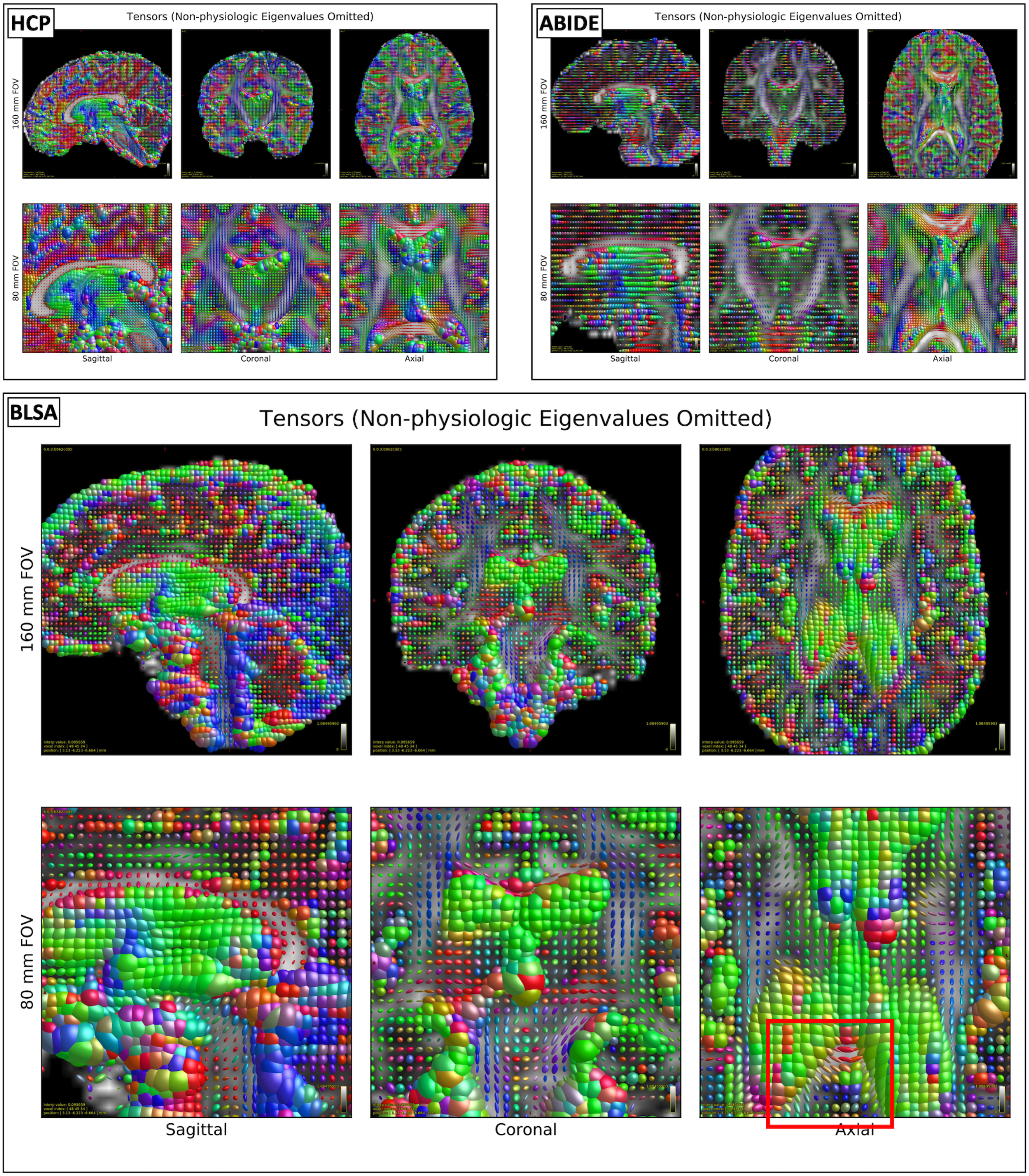


**Supporting Information Figure S8.** QA of tensors. The report renders tensor glyphs. Non-physiologic tensors with eigenvalues larger than 3 times the apparent diffusion coefficient of water at 37°C (>0.009 $mm^2/s$) or less than 0 are omitted, as shown in the axial slices of the ABIDE II session. When interpreting these figures, we verify two aspects. The first is that the tensor fits are reasonable and not mostly omitted or extremely large or small. Since tensors are generally robust, we find unreasonable fits are usually due to mismatching of volumes in the NIFTI files with the gradient schemes in the bvec or bval files. To confirm this, we refer to the FA and MD characterization step (Supporting Information Figure S9). The second is that we also verify the tensors are oriented in physiologically probable directions. Primarily, we identify that tensors along the cortical spinal tracts and corpus callosum in the coronal slices and in the genu and splenium of the corpus callosum in the axial slices follow expected pathways. In line with the gradients plotted in Supporting Information Figure S6 of this supplement, the tensor glyphs for the BLSA session are oriented in a physiologically improbable manner in the splenium of the corpus callosum (red bounding box), suggesting improperly oriented gradients. If we identify errors, we refer to the gradient verification step (Supporting Information Figure S6).

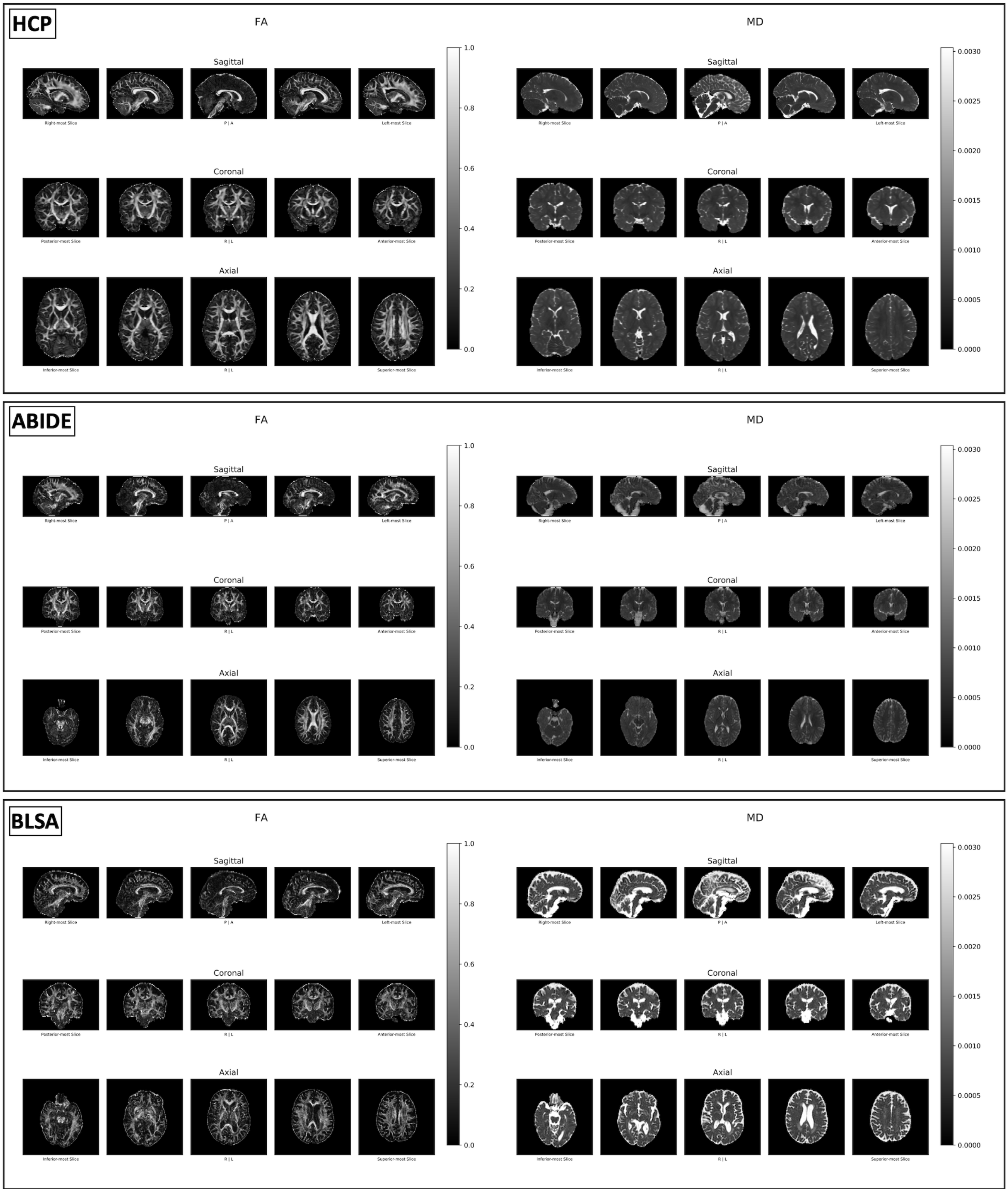


**Supporting Information Figure S9.** QA of FA and MD maps. The report shows five central triplanar slices of the calculated FA and MD maps. When interpreting these plots, we make sure that the FA and MD renderings are within the expected range and follow expected distributions. In other words, we verify that in the FA map white matter appears brighter than gray matter and in the MD map we check that CSF appears bright and the white and gray matter appear homogenous and darker. If we identify errors, we first confirm the tensor visualization also contains errors (Supporting Information Figure S8). The tensor model is generally robust, thus large deviations in fits are usually due to mismatching of volumes in the NIFTI files with the gradient schemes. The first steps we take to fix this error are to verify that the DICOM to NIFTI conversion did not reorder the volumes (Supporting Information Table S1) and that the gradient files were not corrupted otherwise.

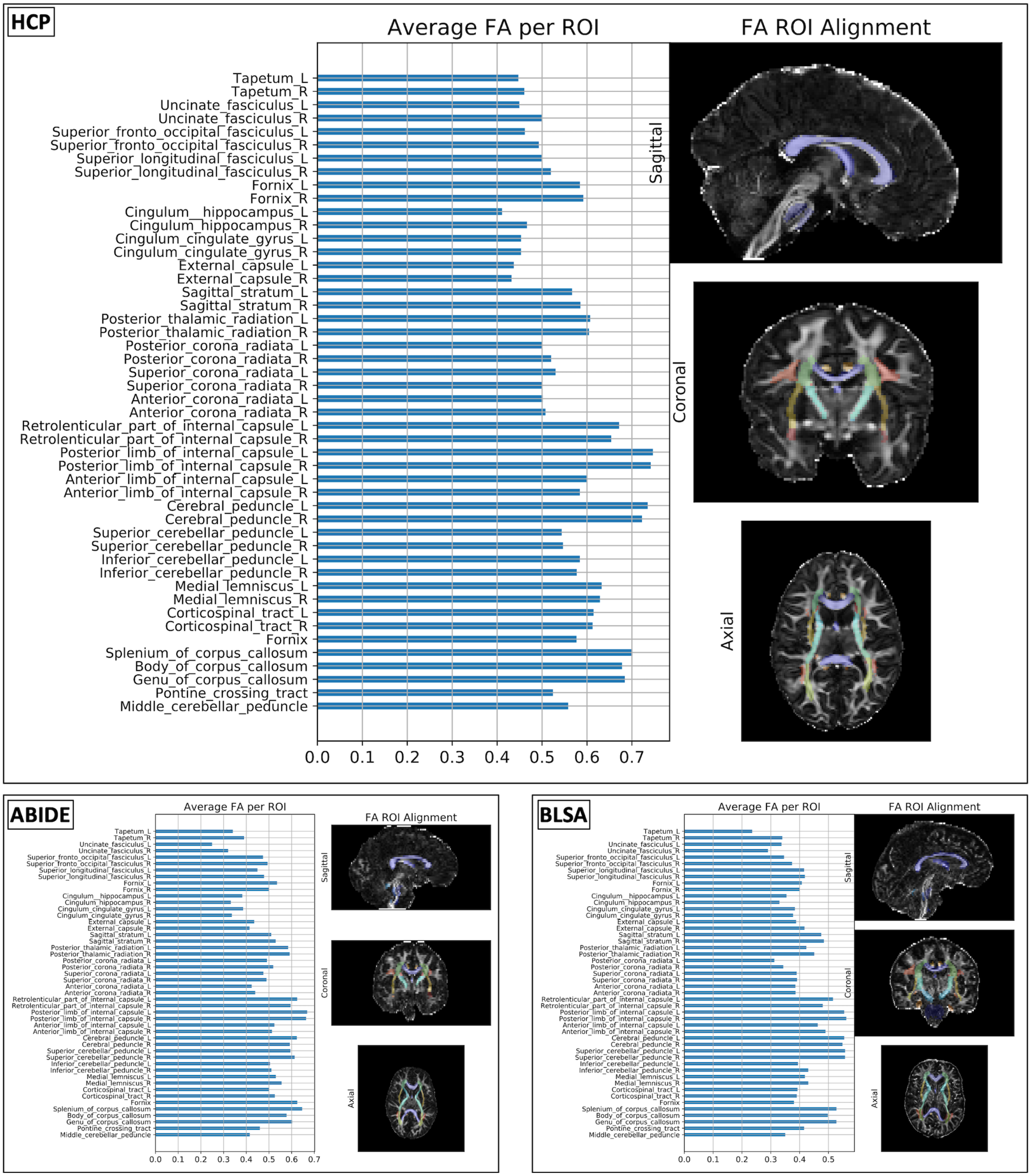


**Supporting Information Figure S10.** QA of the average FA in 48 white matter ROIs. The report shows the average FA of the 48 white matter tracts identified via deformable registration of the Johns Hopkins ICBM DTI 81 white matter atlas to the calculated FA maps. Alignment of the atlas with the FA map is also shown for QA of the registration process. Similar to the FA and MD visualization in Supporting Information Figure S9, we use these plots to understand the quality of tensor fit as a surrogate for quality of the preprocessed DWI output data. Very poor tensor fits often result in either very large or very small regional FA values. We expect proper fits to range from roughly 0.3 to 0.7. If this is not the case, we first verify that the registration was successful using the plots on the right and then we follow the logic outlined for Supporting Information Figure S9.

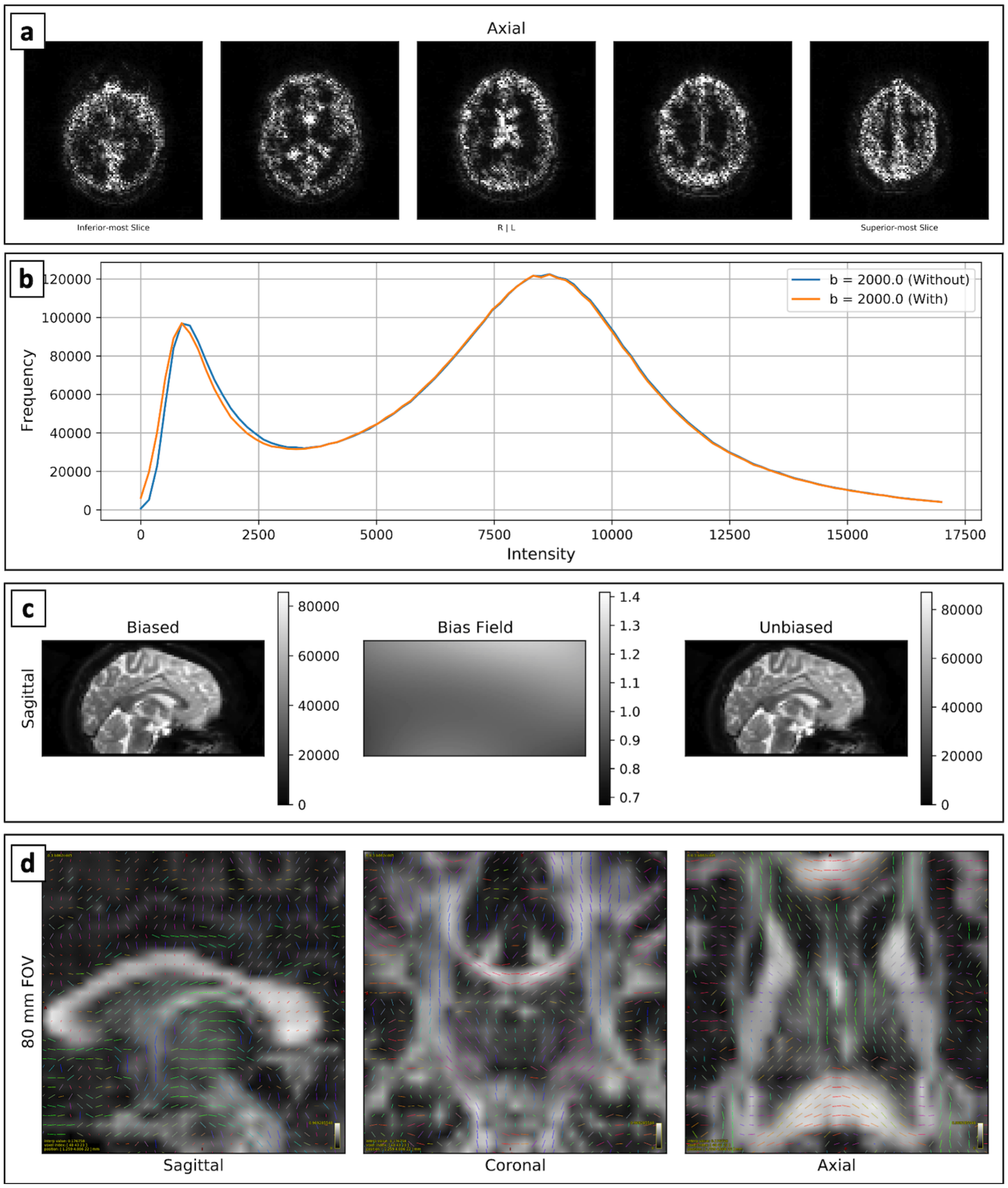


**Supporting Information Figure S11.** Additional Pipeline Options. (a) If the user runs Gibbs de-ringing, the report will display the absolute intensity differences averaged across all b = 0 s/mm$^2$ volumes from before and after correction in triplanar views. Only the axial view is shown here. We expect these differences to be higher at high-contrast interfaces, like between brain parenchyma and CSF as shown. (b) If the user runs Rician correction, the report will display the shell-wise intensity distributions within the brain for each image without and with correction. With the method of moments, we expect to see slight decreases in intensity with correction, as is shown here for the b = 2000 s/mm$^2$ volumes. (c) If the user runs N4 B1 bias field correction, the report will display triplanar views before and after correction as well as the calculated bias field. Only the sagittal view is shown here. The color bars are representative of all intensities in the corresponding volume, and we expect the magnitude of the fields to generally be far smaller than that of the images, as shown here. (d) If the user chooses, the report can also visualize principal eigenvectors instead of tensors. This is especially useful for high in-plane resolution images or up-sampled images where crowding of tensor glyphs can make interpretation difficult. As with the glyphs, these are shown with a 160mm and 80mm FOV. Only the latter is shown here. These visualizations were all taken from the QA report generated on Dataset B.

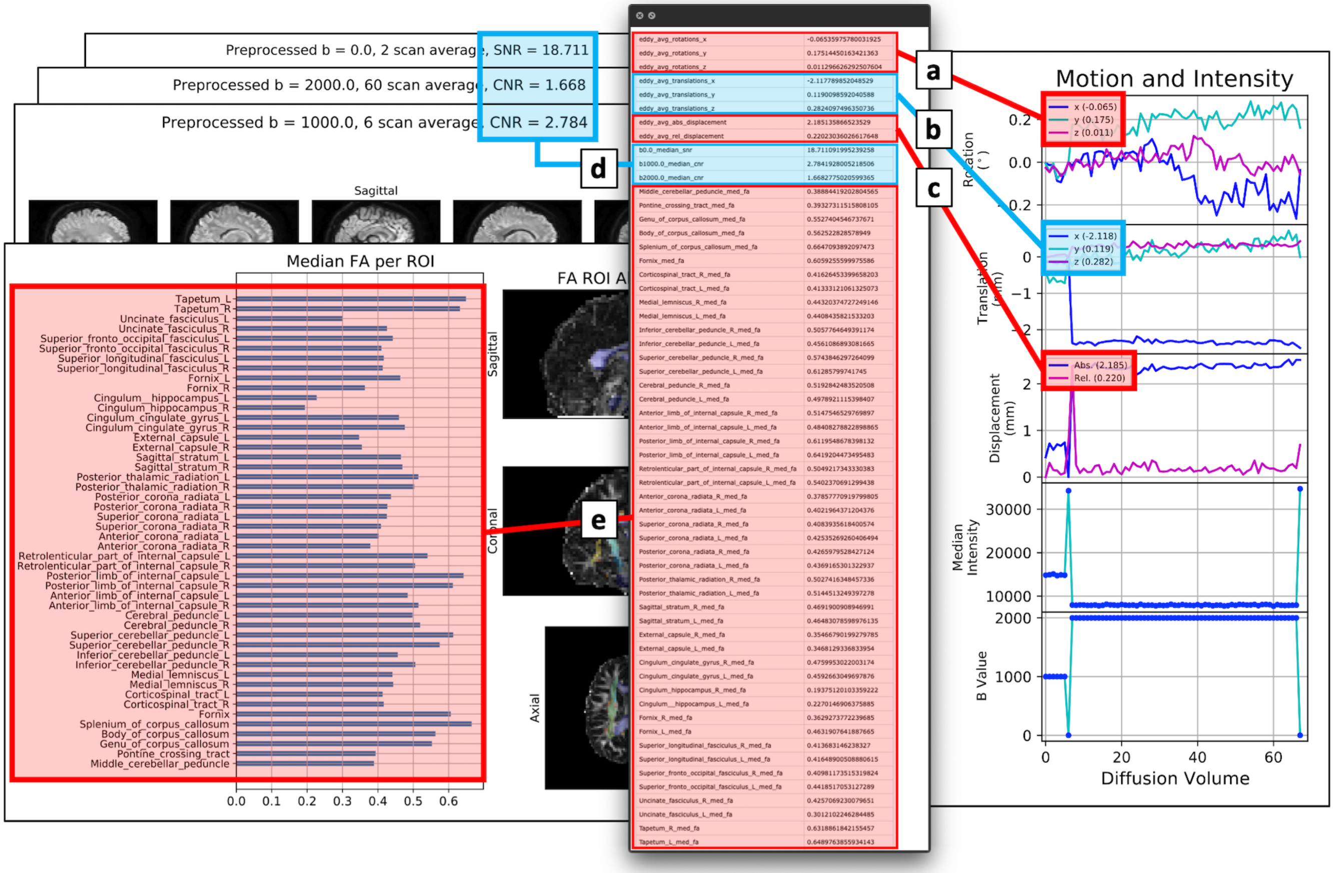


**Supporting Information Figure S12.** Reporting of summary statistics. Summary statistics are reported in the QA document and in an output CSV for user interrogation. The average inter-volume rotation (a), translation (b), and displacement (c) as well as the shell-wise SNR and CNR (d) and regional FA (e) are all tabulated in the CSV. These pages and the CSV information were taken from the QA report generated on Dataset B.

**Supporting Information Table S3.** Comparison of PreQual to the DESIGNER and TORTOISE pipelines.

| | DESIGNER | TORTOISE | PreQual |
|---|---|---|---|
| **Core components** | | | |
| Preprocessing | ✓ | ✓ | ✓ |
| Model fitting | ✓ | ✓ | × |
| Quality assurance | × | × | ✓ |
| **Input handling** | | | |
| Image concatenation (joint processing) | ✓ | ••• | ✓ |
| Inter-image intensity normalization | ✓ | × | ✓ |
| **Preprocessing** | | | |
| B1 bias field correction | ••• | × | ••• |
| CSF-excluded smoothing | ✓ | × | × |
| Eddy current correction | ✓ | ✓ | ✓ |
| Gibbs de-ringing | ✓ | ✓ | ••• |
| Gradient non-linearity correction | × | ✓ | × |
| MP-PCA denoising | ✓ | ✓ | ✓ |
| Motion correction | ✓ | ✓ | ✓ |
| Rician correction | ✓ | × | ••• |
| Slice-wise dropout imputation | ✓ | × | ✓ |
| **Susceptibility correction by input (Dataset B)** | | | |
| **Image 1:** 1 volume b = 0 s/mm² APP; **Image 2:** 60 volume b = 2000 s/mm² APP; **Image 3:** 1 volume b = 0 s/mm² APA; **Image 4:** 6 volume b = 1000 s/mm² APA | | | |
| Images 1 and 2 | × | × | × |
| Images 1, 2, and 3 | ✓ | ✓ | ✓ |
| Images 1, 2, 3, and 4 | ✓[a] | ✓ | ✓ |
| Images 1, 2, and a T1 image | × | ✓ | ✓ |
| Images 1, 2, and a T2 image | × | ✓ | × |
| **Model fitting** | | | |
| DTI | ✓ | ✓ | ✓[b] |
| DKI | ✓ | × | × |
| WMTI | ✓ | × | × |
| MAP-MRI | × | ✓ | × |
| **Quality assurance of…** | | | |
| Each preprocessing step | × | × | ✓ |
| Data before and after preprocessing | × | ✓ | ✓ |
| Gradients and file conversion errors | × | × | ✓ |
| **Software considerations** | | | |
| Containerized | × | × | ✓ |
| Runtime[c] | ~1 hour[d] | ~4 hours[d] | ~1 hour |

✓ Recommended or default on
••• Optional or default off
× Not included

[a] Image 4 not used
[b] Primarily done for QA purposes and not in preparation for secondary analysis
[c] With default parameters and images 1-4 (Dataset B) as input on a 4-core 16GB RAM Ubuntu 18.04 workstation with multithreading
[d] Preprocessing only (no model fitting)

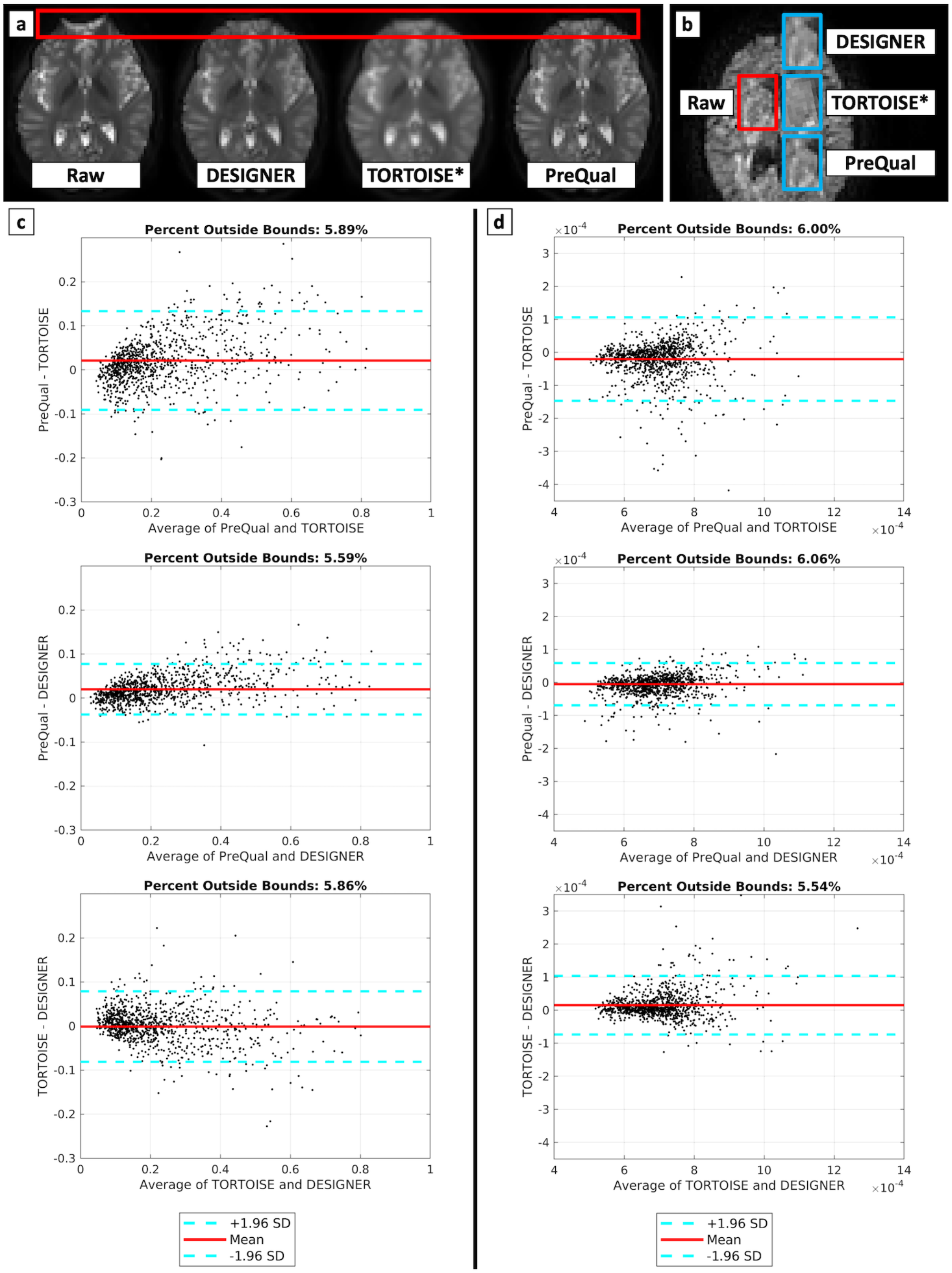


**Supporting Information Figure S13.** Comparison of preprocessed DWI data output by PreQual to those output by the DESIGNER and TORTOISE pipelines on Dataset B. (a) The preprocessed outputs of each pipeline demonstrate correction of susceptibility-induced distortions (red bounding box). (b) The preprocessed outputs of each pipeline (blue bounding boxes) demonstrate decreased noise in a region compared to the raw input (red bounding box) in a diffusion weighted volume. The voxel-wise tensor-based FA (c) and MD (d) scalar values within the brain parenchyma computed by MRTrix3's IWLS estimator on the preprocessed DWI outputs of the three pipelines exhibit good agreement on Bland-Altman plots. Approximately 5-6% of voxels fall outside the 95% confidence intervals. We show 5% of the voxel data points for interpretability, and all images were rigidly registered to the PreQual output to achieve correspondence. All pipelines were run with default or recommended settings (Supporting Information Table S3).

*TORTOISE performs preprocessing in structural MRI space. The TORTOISE output was rigidly registered to the raw DWI image for (a). The output was not registered for (b) to avoid artificial smoothing from interpolation and has close but not exact regional correspondence.
